# Emergence of *Prochlorococcus* in the Tonian oceans and the initiation of Neoproterozoic oxygenation

**DOI:** 10.1101/2023.09.06.556545

**Authors:** Hao Zhang, Sishuo Wang, Tianhua Liao, Sean A. Crowe, Haiwei Luo

## Abstract

*Prochlorococcus* are the smallest and most abundant photosynthetic organisms on Earth, contributing up to 50% of the chlorophyll in the oligotrophic oceans. Despite being important in regulating the carbon cycle in today’s ocean, the ecological significance of *Prochlorococcus* in Earth’s history remains elusive. Our new robustly calibrated molecular clock analysis reveals that *Prochlorococcus* emerged in the deep photic zone of the Tonian (1,000-720 Mya) oceans. The classical light-harvesting antenna complex in Cyanobacteria, i.e., the phycobilisome, was replaced in *Prochlorococcus* by the chlorophyll□based antenna, enabling more efficient use of blue light that penetrates into deeper water. Importantly, *Prochlorococcus* colonization of deep water enhanced access to phosphate, which was abundant in upwelled seawater, but likely scarce in the Tonian surface ocean, promoting expansion of *Prochlorococcus,* displacement of incumbent low-light adapted anoxygenic photoferrotrophs, and associated increases in photosynthetic oxygen production. Colonization of deeper waters would also have improved access to ammonium, leading to the neutral loss of nitrate utilization genes. Our research thus documents the conspicuous emergence of new photosynthetic bacterial lineages in the run-up to the Neoproterozoic oxygenation event, implying an additional layer of eco-evolutionary complexity during this pivotal interval in Earth’s history.

## Introduction

*Prochlorococcus* host more than half of the chlorophyll biomass in oligotrophic oceans (Partensky and Garczarek, 2010). As key primary producers in the modern ocean, *Prochlorococcus* fix as much as four gigatons of carbon each year and are the foundation of the marine carbon cycle and food web (Flombaum et al., 2013). The success of *Prochlorococcus* in today’s oceans has been attributed to multiple physiological features. Importantly, *Prochlorococcus* use divinyl chlorophyll (DVChl) for harnessing light energy (Ralf and Repeta, 1992). DVChl harvests blue light much more efficiently than the more common monovinyl chlorophyll used by other cyanobacterial lineages and it thus enables *Prochlorococcus* to thrive in the deepest layers of the euphotic zone, where blue light dominates (Ito and Tanaka, 2011). *Prochlorococcus* is also the smallest photosynthetic organism on Earth (Partensky et al., 1999). As a result, their high surface-to-volume ratio provides enhanced nutrient acquisition efficiency, which together with effective blue-light absorption promotes photosynthesis by *Prochlorococcus* in oligotrophic tropical and subtropical oceans (Partensky et al., 1999). In today’s ocean, *Prochlorococcus* is also the dominant phototroph in more nutrient-rich, oxygen-depleted anoxic marine zones (AMZ) in the eastern tropical North and South Pacific Oceans (Goericke et al., 2000; Lavin et al., 2010). Since the AMZ lineages represent the earliest-split branches of *Prochlorococcus*, it has been proposed that *Prochlorococcus* emerged in low-oxygen environments and, by extension, contributed to early ocean oxygenation (Ulloa et al., 2021).

The emergence of an early branch of *Prochlorococcus* (named SBE-LCA) during the Cryogenian based on our recent study (Zhang et al. 2021) implies the origin of the total *Prochlorococcus* group earlier in the Proterozoic Eon. Such an earlier origin then suggests that *Prochlorococcus* might have contributed to the cyanobacterial dominated primary production that supported marine biogeochemical cycles and underpinned dynamic ocean chemistry in the run-up to the Sturtian Snowball Earth glaciation. This was an important period in Earth’s history that ultimately led to an Earth system succession and the rise of eukaryotes with algae emerging as key primary producers before the end of the Cryogenian (Brocks et al. 2017). Deciphering the potential role, if any, that *Prochlorococcus* might have played depends critically on an accurate estimate of the origin time for *Prochlorococcus*.

By far, however, the divergence time of *Prochlorococcus* remains contentious from ∼200 Mya to ∼1,000 Mya (Sánchez-Baracaldo et al., 2014; Sánchez-Baracaldo, 2015; Schirrmeister et al., 2015; Sánchez-Baracaldo et al., 2017; Boden et al., 2021; Fournier et al., 2021; Martinez-Gutierrez et al., 2023). The discrepancies in *Prochlorococcus* divergence times are likely caused by variable use of fossils and other time calibrations and the use of alternative gene sets, clock models, and tree topologies (Warnock et al., 2012; Duchêne et al., 2014; dos Reis et al., 2015) (see Supplementary Discussion). Despite their importance, these factors were rarely tested rigorously thus making it difficult to evaluate the accuracy of age estimates in previous studies. Importantly, even if all these factors are well tested and controlled, the rarity of cyanobacterial fossils poses a notable challenge in determining the antiquity of *Prochlorococcus*, particularly considering the lack of maximum age information when only cyanobacterial fossils were used (Zhang et al., 2021). Indeed, applying maximum age constraints at the calibration nodes strongly impacts posterior age estimates of uncalibrated lineages (Hedges et al., 2018; Morris et al., 2018; Wang and Luo, 2021). However, informative maximum age constraints are typically missing from microbial fossils and can only be found in some animal and plant fossils. Moreover, the recent availability of several genome sequences of uncultivated basal *Prochlorococcus* lineages (Ulloa et al., 2021) requires re-estimation of the antiquity of *Prochlorococcus*, which should also include an evaluation of the factors that influence the accuracy of posterior ages.

Here, we leverage the abundant plant and algal fossils and the well-established plant plastid endosymbiosis theory to develop a new pipeline that systematically tests the factors that influence the accuracy of posterior ages and thus yields robust estimates of *Prochlorococcus* antiquity. The plant plastid endosymbiosis theory states that the origin of all plastids in photosynthetic eukaryotes, except for the photosynthetic amoeba *Paulinella* (Marin et al., 2005), can be traced back to an ancient primary endosymbiosis involving a eukaryote and a cyanobacterium (Gray, 1992; Bhattacharya and Medlin, 1995; Keeling, 2013; Ponce-Toledo et al., 2017). Plant plastid endosymbiosis theory thus ties the evolutionary histories of cyanobacteria to those of photosynthetic eukaryotes. Eukaryotic fossils are indeed being increasingly used in dating the evolution of Cyanobacteria (Shih et al., 2016; Sánchez-Baracaldo et al., 2017; Fournier et al., 2021). Our strategy builds on these by: 1) using more fossil-based age constraints (including maximum age constraints) on eukaryotic lineages with more complete taxonomic sampling including lineages that have undergone secondary endosymbiosis; and 2) applying a Bayesian sequential dating approach to better constrain the divergence time of eukaryotic lineages thereby propagating more accurate time information to Cyanobacteria, including *Prochlorococcus*. Note that the Bayesian sequential method used here is different from the commonly used secondary dating method (using time estimates from previous studies as calibrations) (Heckman et al., 2001; Aoki et al., 2013; Chriki-Adeeb and Chriki, 2016) for two reasons. First, the time prior used in sequential dating analysis follows a specific probability distribution, but secondary calibrations ignore the uncertainties on age estimates. Second, the sequential dating analysis contains two steps each based on a distinct set of molecular data, while secondary dating analysis largely relies on the same molecular dataset (gene alignments) (dos Reis et al., 2018). These joint analyses led us to conclude that *Prochlorococcus* arose in the Tonian (1,000-720 Mya) oceans. Further, population genetic analysis implies that *Prochlorococcus* was born through a founder effect, and this strengthens the idea that *Prochlorococcus* emerged in the deep photic zone, a unique niche separated from upper waters where its ancestors (i.e., the last common ancestor of *Prochlorococcus* and its sister clade in the genus *Synechococcus*) thrived.

## Results and Discussion

### *Prochlorococcus* originated in the Tonian ocean

We implemented a Bayesian sequential dating method that takes advantage of the abundant fossil records (Fig. 1a; Fig. S1a) available from photosynthetic eukaryotes to calibrate the evolution of Cyanobacteria. In our implementations, the posterior ages of eukaryotic lineages derived from the first-step of the sequential analysis were used as the time priors to calibrate Cyanobacteria evolution where only a few time constraints are available. To achieve this goal, we first implemented the genome-scale dating analysis of the eukaryotic lineages and confirmed that the posterior ages of the crown eukaryotic group were consistent with the previous estimates at ∼1.6 Gya (Parfrey et al., 2011; Betts et al., 2018; Wang and Luo, 2021) (Fig. S1b). We then compared the posterior age distributions of eukaryotic nodes from the first-step analysis to the distributions of the effective time prior on the corresponding nodes in the second-step analysis. The nearly identical distributions found in the comparisons suggest that the Bayesian sequential dating approach works well in these cases (Fig. S2).

**Fig. 1.**
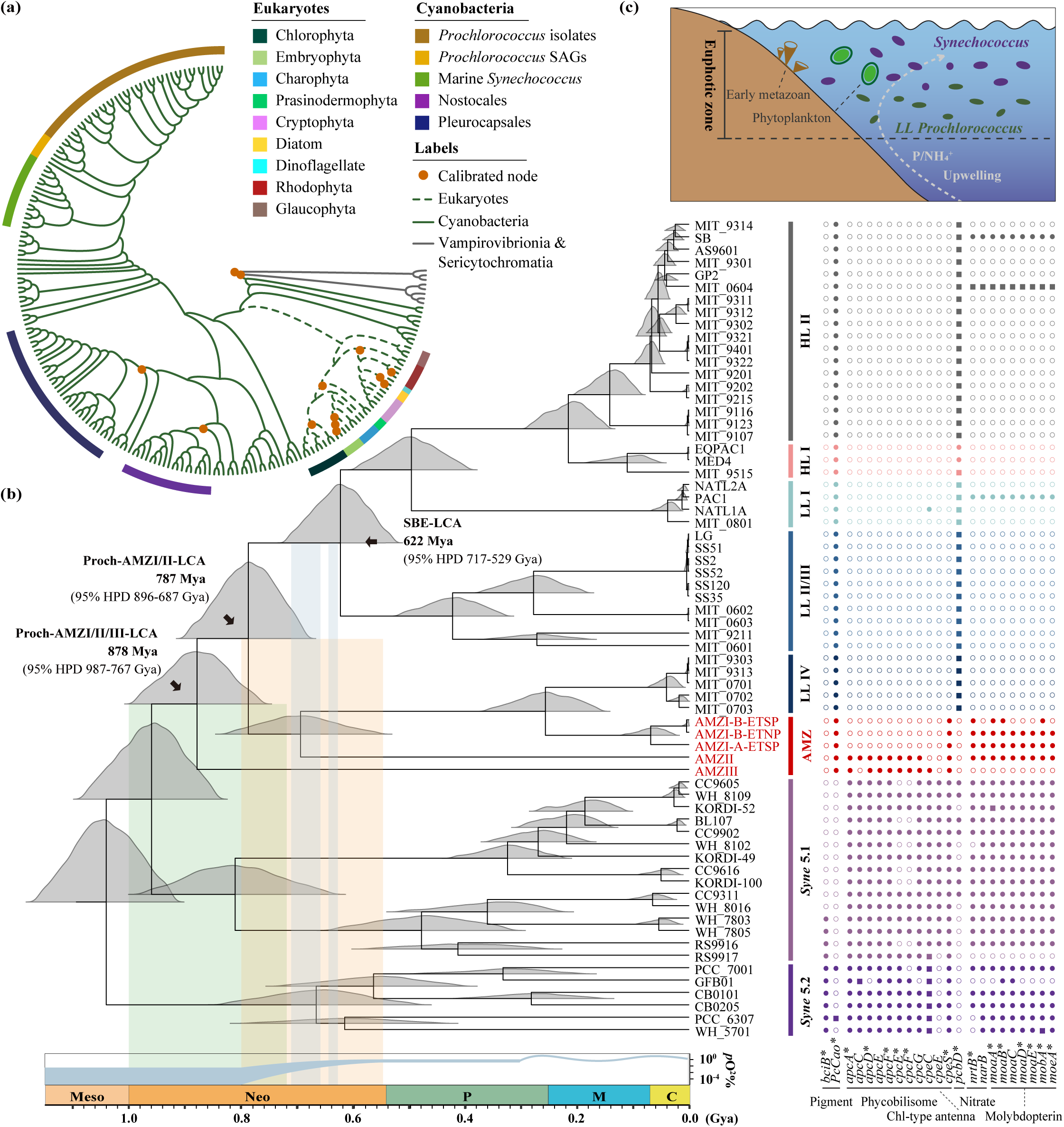
The evolutionary timeline of *Prochlorococcus* estimated with plastid-based strategy. (a) The diagram shows the phylogenetic relations among Cyanobacteria and eukaryotic species. The green solid line, black solid line and green dashed line in the cladogram leading to the tip of oxygenic Cyanobacteria, anoxygenic Vampirovibrionia and Sericytochromatia and eukaryotic species, respectively. The calibrated nodes in molecular clock analysis are marked with orange circle (see calibration justifications in Supplementary Methods). (b) (left) The *Prochlorococcus* evolutionary timeline estimated with the focal molecular dating strategy using Bayesian sequential dating approach based on the 5-partition eukaryotic genome-scale data (in the first step dating analysis) and fully partitioned 12-gene dataset “T30” (in the second step dating analysis) under independent rate clock model. The posterior age distribution is provided next to the ancestral node. The atmospheric oxygen level at the geological time scale is adapted from Lyons et al., 2014, which is represented by the percent of present atmospheric oxygen level (PAL). The vertical bars with green, orange and blue colors represent the time of Tonian, the time of NOE, and the time of Sturtian (left) and Marinoan (right) glaciation, respectively. (right) Phyletic pattern of key gene families. Solid square, solid circle and open circle next to each analyzed genome represent multi-copy gene family, single-copy gene family and absence of the gene family, respectively, in the extant genomes. Gene families marked with asterisk (*) were consistently estimated to be gained or lost by using both AnGST and GeneRax. (c) Diagram illustrating the biogeochemical environments when *Prochlorococcus* arose from its *Synechococcus* ancestor.

Our dating analysis implies that the last common ancestor (LCA) of *Prochlorococcus* (denoted as “Proch-AMZI/II/III-LCA”) emerged within the Tonian period at 878 Mya (95% HPD: 987-767 Mya) (Fig. 1b). Meanwhile, the LCA of *Prochlorococcus* clades HL, LL and AMZI/II (denoted as “Proch-AMZI/II-LCA”) evolved at 787 Mya (95% HPD: 896-687 Mya), which coincided with the early stage of the Neoproterozoic Oxygenation Event (NOE; 800-550 Mya) (Fig. 1b). In this case, the branch leading to the LCA of *Prochlorococcus* HL, LLI, and LLII/III clades (denoted as “SBE-LCA” to keep consistency with (Zhang et al., 2021)) spanned the time that encompassed the duration of the Neoproterozoic Snowball Earth events (645-635 Mya for Marinoan glaciation and 717-659 Mya for Sturtian glaciation), supporting the main conclusion of the previous study, which used the traditional cyanobacterial fossil-based strategy (Zhang et al., 2021).

To validate the evolutionary timeline of *Prochlorococcus*, we performed a series of tests by using different molecular data, clock models, fossil calibrations and species tree topologies. The posterior ages estimated with all these alternative settings are largely consistent with that estimated with the focal strategy (detailed above) and fully support the origin of *Prochlorococcus* (Proch-AMZI/II/III-LCA) in the Tonian period (1,000-720 Mya), even when the maximum root age changed from 3.8 Gya to 4.5 Gya (Fig. S3). Moreover, the emergence of Proch-AMZI/II-LCA was consistently estimated to occur in the early stages of the NOE (800-550 Mya) (Fig. S4), and that the branch leading to the SBE-LCA encompasses the Snowball Earth events in all the dating analyses (Fig. S5; see Supplementary Results).

### The emergence of *Prochlorococcus* is associated with a founder effect

The method for inferring selection efficiency in deep time was developed (Zhang, 2000) and recently improved (Luo et al., 2017), which involves nonsynonymous substitutions only and compares the rate of nonsynonymous substitutions leading to replacements of physicochemically dissimilar amino acids (i.e., radical changes; *d_R_*) to the rate of nonsynonymous substitutions leading to replacements of physicochemically similar amino acids (i.e., conservative changes; *d_C_*). Since radical changes are more likely to be deleterious than conservative changes (E. Zuckerkandl, 1965; Dayhoff, 1972), an excess of the radical changes in a deeply branching lineage compared to its sister lineage suggests random fixation of deleterious mutations by genetic drift in the former. Using this method, we found a significant increase of the *d*_R_/*d*_C_ ratio across the genomic regions in the branch leading to Proch-AMZI/II/III-LCA relative to the branch leading to the LCA of *Synechococcus* clade 5.1 (Fig. 2a), indicating that the emergence of *Prochlorococcus* was accompanied by a significant reduction of the efficiency of purifying selection and a potentially severe reduction in the effective population size (*N_e_*), allowing for an accelerated accumulation of deleterious mutations through genetic drift.

**Fig. 2.**
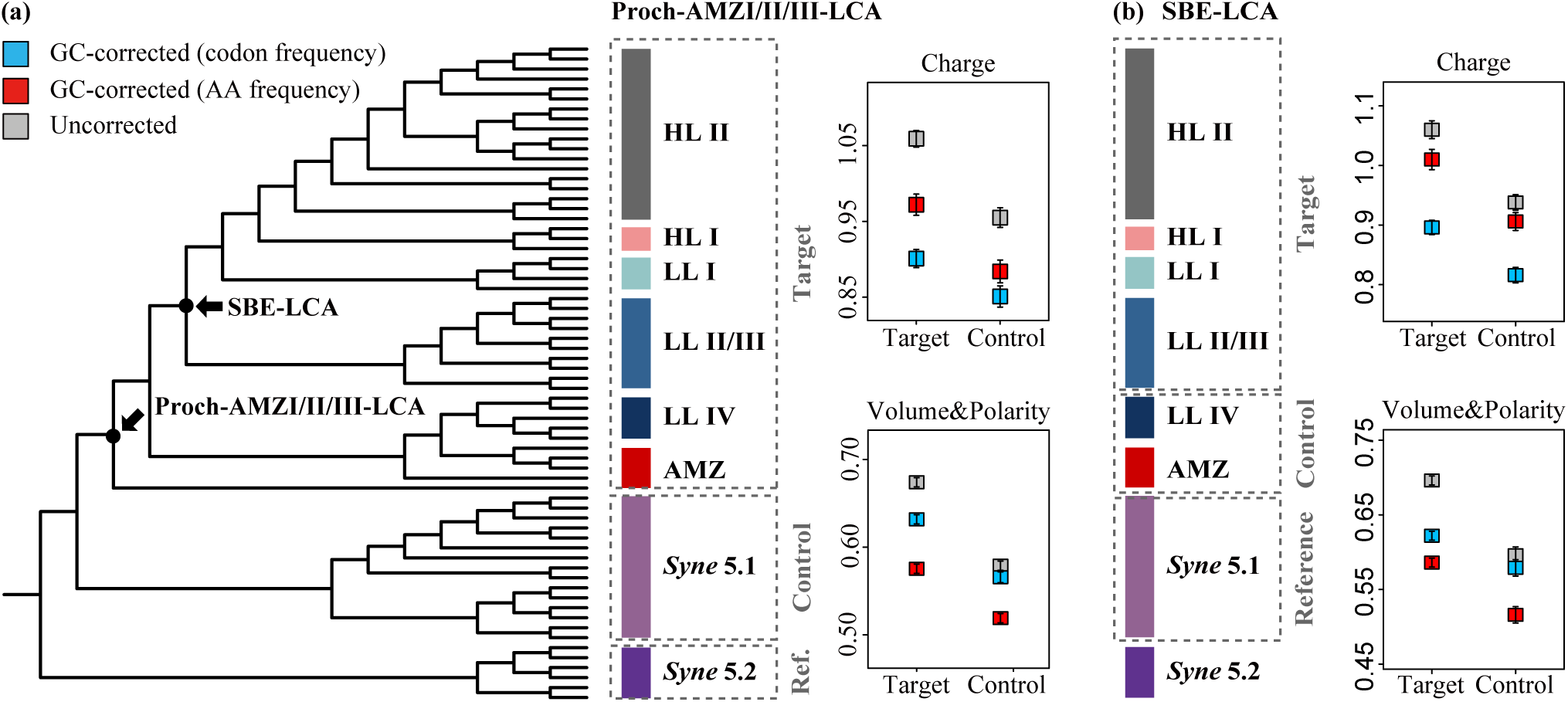
Diagram illustrating the schemes and the results of *d*_R_/*d*_C_ comparison. The genome-wide means of *d*_R_/*d*_C_ values at the ancestral branches (‘Target’) leading to (a) Proch-AMZI/II/III-LCA and (b) SBE-LCA compared with that at the sister lineages (‘Control’). The *d*_R_/*d*_C_ values were classified based on the physicochemical classification of the amino acids by charge or by volume and polarity, and were either GC-corrected by codon frequency (blue), GC-corrected by amino acid (AA) frequency (red) or uncorrected (gray). Error bars of *d*_R_/*d*_C_ represent the standard error of the mean.

We propose that the colonization of the deep photic zone by *Prochlorococcus* started by a seed population that obtained the ability to use divinyl chlorophylls. Therefore, the reduced *N_e_* upon the emergence of *Prochlorococcus* was likely caused by a “founder effect”, which suggests that a new population was established by a few individuals from a larger ancestral population (Barton et al., 1984). Note that this mechanism is different from population bottleneck, which is often caused by the environmental disasters. The latter has been used to explain the reduced *N_e_* occurring on the branch leading to SBE-LCA (Zhang et al., 2021). That conclusion is confirmed here with the inclusion of new SAG genomes, as the *d*_R_/*d*_C_ values in the branch leading to SBE-LCA were significantly elevated compared to the those in the branches leading to *Prochlorococcus* LLIV and AMZ clades (Fig. 2b)

### *Prochlorococcus* genomic changes and niche adaptation

As the only phytoplankton group using divinyl chlorophyll (DVChl) for harvesting light energy (Ralf and Repeta, 1992), *Prochlorococcus* lost the gene *bciB* that performs the conversion of DVChl to MVChl (monovinyl chlorophyll) at Proch-AMZI/II/III-LCA, thereby promoting the accumulation of DVChl in their membranes (Ito et al., 2008). Compared to the MVChl used by *Synechococcus*, DVChl more efficiently absorbs blue light that penetrates to deeper waters than other photosynthetically active radiation (Ito and Tanaka, 2011). Likewise, the gain of the *PcCao* gene for the synthesis of chlorophyll b (Satoh and Tanaka, 2006) in Proch-AMZI/II/III-LCA also enables *Prochlorococcus* ancestors to efficiently harvest blue light at exceedingly low intensities characteristic of deep waters (Hess et al., 2001) (Fig. 1b). These genomic changes would have enabled *Prochlorococcus* ancestors to explore the deep photic zone, a niche where its *Synechococcus* ancestor would not thrive.

We infer that dwelling in deeper waters confers competitive advantages upon *Prochlorococcus*. A well-known advantage emerges under conditions of strong phosphorus (P) limitation, whereby deep-dwelling low-light adapted phototrophs gain first access to upwelling phosphorus (Jones et al., 2015; Ozaki et al., 2019). The low-light advantage conferred on the earliest *Prochlorococcus* cells through the adoption of divinyl chlorophyll would have enhanced their access to P, relative to contemporary algae and its own ancestors. It would also have increased access to NH_4_^+^ upwelling from deeper anoxic waters obviating the need for nitrate assimilation (Michiels et al., 2017) (Fig. 1c).

Unlike most cyanobacteria, which use the phycobilisome as the photosynthetic antenna, the main light-harvesting antenna of *Prochlorococcus* is made up of prochlorophyte chlorophyll-binding (Pcb) protein (Biller et al., 2015). By reconstructing the gene evolutionary paths with *Prochlorococcus* SAGs included, we found that the phycobilisome genes (*apcACDEF*, *cpcEFG*, and *cpeCES*) were present in both Proch-AMZI/II/III-LCA and Proch-AMZI/II-LCA and lost at SBE-LCA. Using the same approach, we found that the cholorophyll-binding protein (encoded by *pcbD*) was obtained at Proch-AMZI/II-LCA. The replacement of the photosynthetic antenna thus did not co-occur with the emergence of *Prochlorococcus*. We note that gene absence in SAGs could result from the incomplete nature of the SAG genomes, however, in this case, simultaneous absence of the *pcbD* gene in all the five SAGs that are more than 80% complete seems unlikely.

The phycobilisome constitutes as much as 50%-60% of the soluble proteins in *Synechococcus* (Grossman et al., 1993). The loss of the phycobilisome was thought to reduce nitrogen (N) investments by at least 40% in *Prochlorococcus* (Ting et al., 2002). Therefore, the losses of phycobilisome genes was once considered to be favored by *Prochlorococcus* conferred advantages to inhabiting oligotrophic oceans (Ting et al., 2002). However, since the colonization of *Prochlorococcus* in the deep photic zone of the Tonian ocean conferred them advantages in acquiring the limiting nutrients like ammonium and phosphate upwelled from the deep ocean (Jones et al., 2015; Michiels et al., 2017; Ozaki et al., 2019), the loss of the phycobilisome in *Prochlorococcus* was unlikely driven by selection for metabolic efficiency. Instead, as the chlorophyll□based light□harvesting complex gradually became the primary photosynthetic antenna in *Prochlorococcus*, the phycobilisome genes were more likely subject to relaxed purifying selection and thus neutral losses.

Despite the absence of the nitrate utilization genes in most *Prochlorococcus* isolates, some uncultivated *Prochlorococcus* lineages contain these genes (Martiny et al., 2009; Berube et al., 2015; Berube et al., 2019). By including *Prochlorococcus* SAGs that contain the assimilatory nitrate transporter gene (*nrtB*) and the assimilatory nitrate reductase gene (*narB*) in our reconstructions, we inferred the presence of both genes in Proch-AMZI/II/III-LCA and Proch-AMZI/II-LCA and the losses of these genes at SBE-LCA (Fig. 1b). The absence of nitrate utilization genes (*narB* and *nrtB*) in cultured *Prochlorococcus* strains has been attributed to biased taxon sampling, since the *Prochlorococcus* were primarily cultured from ocean regions where P instead of N is the primarily limiting nutrient (Berube et al., 2015). However, phylogenetic analysis that included both *Prochlorococcus* and *Synechococcus* showed topological congruence of the *narB* gene tree with the species tree, suggesting vertical *narB* gene inheritance (Berube et al., 2019). By including *Prochlorococcus* SAGs that contain the assimilatory nitrate transporter gene (*nrtB*) and the assimilatory nitrate reductase gene (*narB*) in our reconstructions, we inferred the presence of both genes in Proch-AMZI/II/III-LCA and Proch-AMZI/II-LCA and the losses of these genes at SBE-LCA (Fig. 1b). The mechanism underlying the loss of nitrate utilization genes in *Prochlorococcus* was once attributed to relaxed selection efficiency due to the low level of nitrate caused by intensive denitrification and anammox (anaerobic ammonium oxidation) activity (Canfield et al., 2008; Johnston et al., 2009). However, since *Prochlorococcus* originated in the deep photic zone where ammonium would likely have been supplied through upwelling (Fig. 1c) (Michiels et al., 2017), the loss of the nitrate utilization genes in early *Prochlorococcus* is more likely to be the result of a switch of its N source from nitrate to ammonium, which was again a neutral process.

In addition to the nitrate utilization genes, we inferred the losses of molybdopterin biosynthesis genes at SBE-LCA. Since these genes are known to co-locate with the nitrate reductase genes in *Synechococcus* (Rubio et al., 1998; Palenik et al., 2003), they may function as the cofactor of nitrate reductase in *Prochlorococcus*. Therefore, the loss of nitrate reductase in *Prochlorococcus* likely rendered the molybdopterin dispensable and thus led to the losses of molybdopterin biosynthesis genes (*moaABCDE*, *mobA* and *moeA*) at SBE-LCA.

### Geochemical context that supports the emergence of *Prochlorococcus* in Tonian ocean

A Tonian, pre-Sturtian, age for the emergence of the *Prochlorococcus* lineage would have been set against the backdrop of dynamic ocean chemistry characterized by a variably oxygenated surface ocean that was co-populated by diverse microbial eukaryotes, including phytoplankton and the earliest metazoans (Erwin et al., 2011) (Fig. 1c). Multi-proxy data, collected from geographically diverse sites, converge on Tonian deep oceans that were pervasively anoxic across 60-80% of the ocean floor, or more (Tahata et al., 2015; Lau et al., 2017). The deep anoxic oceans were predominantly ferruginous in nature, albeit with evidence for transient, spatially restricted euxinia, and possibly even brief (<0.5 Myr) episodes of pervasive deep ocean oxygenation (Stolper and Keller, 2018; Tostevin and Mills, 2020). Surface waters, by contrast, were modestly, though perhaps increasingly, oxygenated across the Tonian with evidence for the episodic and persistent intrusion of anoxic deep waters into the surface oceans (Zhang et al., 2022; Clarkson et al., 2023), with corresponding implications for nutrient cycling and availability and for marine life in the euphotic zone.

In this way, the emergence of the *Prochlorococcus* would have restricted the flux of nutrients to surface waters and limited the contributions of higher-light adapted phytoplankton like algae to primary production. This is supported through the biomarker record, which indicates a dominance of cyanobacterial primary production until after the Sturtian glaciation (Brocks, 2018). It is also supported by the Tonian N-isotope record, which implies a limited contribution from nitrate assimilation to primary production at this time (Kang et al., 2023). Importantly, the adaptation of the *Prochlorococcus* LCA to lower-light would also allow it to better compete with low-light adapted anoxygenic phototrophs that would likely have populated anoxic regions of the Tonian euphotic zone with strong potential to cause ocean oxygenation (Johnston et al., 2009; Jones et al., 2015; Ozaki et al., 2019). Competition between oxygenic photosynthetic cyanobacteria and anoxygenic phototrophs that oxidize ferrous iron (photoferrotophs), is known to limit photosynthetic oxygen production (Jones et al., 2015; Ozaki et al., 2019). Whereas the accumulation of O_2_ produced through photosynthesis ultimately depends on organic matter burial (Berner, 1991), oxygen production fluxes through organic matter burial scale with the fraction of total photosynthetic production that is oxygenic rather than anoxygenic (Johnston et al., 2009; Ozaki et al., 2019). In this way, the capacity of the *Prochlorococcus* LCA to grow in deeper waters would have enhanced its ability to access upwelling phosphorus thereby increasing the oxygenic fraction of total photosynthesis with potential to initiate a positive feedback on oxygenation (Ozaki et al., 2019).

### Implications for Neoproterozoic biogeochemical cycling

In accordance with previous research (Ulloa et al., 2021), our analysis suggests that *Prochlorococcus* diverged before transitioning from the use of the phycobilisome to the chlorophyll□based light□harvesting complex. Therefore, while the divinyl pigment synthesis gene was obtained at the earliest *Prochlorococcus* (i.e., Proch-AMZI/II/III-LCA) to enable their exploration of the deep photic zone, our analyses imply that *Prochlorococcus* would not have strongly influenced the Neoproterozoic Earth system until the emergence of Proch-AMZI/II-LCA lineage during which the photosynthetic antenna was replaced. Our dating analysis shows that the emergence of this lineage indeed broadly coincides with geochemical signals for the early stages of Neoproterozoic Oxygenation Event (NOE).

The NOE is widely considered the second stage of Earth’s protracted oxygenation history, during which atmospheric and marine oxygen concentrations rose to levels exceeding those that characterized the proceeding mid-Proterozoic (Och and Shields-Zhou, 2012; Lyons et al., 2014). This rise in atmospheric oxygen ultimately promoted the emergence of large and complex animal life (Knoll and Nowak, 2017). The initiation of the NOE was previously linked to eukaryotic algae (Lyons et al., 2014; Brocks et al., 2017) or marine picocyanobacteria (Sánchez-Baracaldo et al., 2014; Braakman et al., 2017). Our results show support for the latter assumption and indicate that early *Prochlorococcus* may have played a role in enhancing oxygen production, with potential to initiate the NOE, which was likely ultimately accelerated through increasing efficiency in the biological carbon pump driven by a progressively a larger role for eukaryotes. Such an elevated role for eukaryotes is clearly marked in the geologic record through an increase in sterane-hopane ratios following the Sturtian glaciation (Brocks et al., 2017). Pre-Sturtian, Tonian oxygenation, was thus likely driven by bacterially dominated carbon cycling, the timing of which strongly coincides with the emergence of low-light adapted *Prochlorococcus*, their ensuing colonization of the deep photic zone, and capacity to displace incumbent anoxygenic photoferrotrophs thereby increasing oxygen production. This hypothesis could be further tested through geochemical analyses that help refine estimates for Tonian ocean oxygen contents and through biogeochemical models that can quantify the potential effects of *Prochlorococcus* emergence on the global carbon and oxygen cycles. Note that the inference of the roles of *Prochlorococcus* in facilitating Neoproterozoic biogeochemical cycle relies on the accurate molecular dating analysis. Given the heavy debate about the antiquity of Cyanobacteria, including *Prochlorococcus*, we have extensively discussed the previous dating analyses and pointed out their methodological deficiencies (see details in Supplementary Discussion).

### Concluding remarks

We have integrated molecular clocks, evolutionary genetic analyses, and comparative genomics along with knowledge of paleo-geochemistry to reconstruct the early evolution of *Prochlorococcus* and infer their possible impact on the evolution of the Neoproterozoic Earth system. The abundant fossil calibrations ‘borrowed’ from photosynthetic eukaryotes and the new and robust molecular dating pipeline pinpoint the origin of *Prochlorococcus* to the Tonian oceans. Comparative genomics suggests that gains and losses of a few important genes conferred the early ancestral *Prochlorococcus* with an unprecedented ability to absorb the blue light that effectively penetrates water, empowering early *Prochlorococcus* to colonize the deep photic zone, a niche distinctly different from the better illuminated surface waters supporting other phytoplankton groups including its *Synechococcus* ancestor and early Eukaryotes. The use of the evolutionary genetic proxy, *d*_R_/*d*_C_, implies that the earliest *Prochlorococcus* had a highly reduced effective population size compared to its *Synechococcus* ancestor, likely reflecting a founder effect and strengthening the idea that the deep photic zone was an ecological niche uniquely accessible to the earliest *Prochlorococcus*. This niche differentiation allowed the earliest *Prochlorococcus* to avoid fierce competition with its *Synechococcus* ancestor and other phytoplankton groups and gave them easy access to phosphate that otherwise strongly limited primary productivity in the Tonian oceans. While all of these co-existing planktonic groups are likely to have contributed to the NOE, the emergence of *Prochlorococcus* in a previously uncolonized habitat with a unique light regime and a higher phosphate accessibility, followed by the acquisition of a more efficient light-harvesting system, may have facilitated a potentially rapid population expansion, which ultimately enhanced the oxygen production in Tonian oceans and facilitated the initiation of the Neoproterozoic Oxygenation Event.

## Materials and Methods

The nuclear-encoded and plastid-encoded protein sequences of eukaryotic representatives were obtained from Dicots PLAZA (5.0) database and NCBI RefSeq release of plastid database and were used for the first-step and the second-step Bayesian sequential dating analysis, respectively. The genomic sequences of Cyanobacteria (including *Prochlorococcus* SAGs), Vampirovibrionia and Sericytochromatia were also used for the second-step sequential dating analysis, which were obtained from GenBank and Integrated Microbial Genomes (IMG) database (Supplementary Data 1).

For the first-step sequential dating analysis, the orthologous gene families shared by eukaryotic species were identified by searching against a pre-compiled eukaryotic nuclear gene dataset (Strassert et al., 2021). Since genome-scale dating of eukaryotic lineages is very slow with fully partitioned molecular data and is prone to have reduced coverage probability (the probability that the 95% HPD interval of posterior ages contains the true age) due to the large number of partitions (Angelis et al., 2018), we clustered the eukaryotic gene families into 1-, 3-, 5-, 7-, and 9-partitions by using Gaussian Mixture Model (GMM) clustering method based on their estimated evolutionary rates (see details in Supplementary Methods). The 5-partition data was finally adopted for the first-step dating analysis because of the lowest Akaike Information Criterion (AIC) value (Fig. S1b) and the high similarity between the derived distributions of posterior ages and effective time prior (Fig. S2). The eukaryotic fossil-based calibrations used in this first-step dating analysis were adapted from published studies (Fig. S1a; Table S1; See details in Supplementary Methods).

For the second-step sequential dating analysis, the orthologous gene families were firstly identified by searching against a pre-compiled plastid marker gene dataset (Ponce-Toledo et al., 2017) and then sorted by ΔLL (i.e., the difference in log-likelihood values of the gene tree constructed with and without the backbone species tree topology) and evolutionary rate difference (Supplementary Data 2) (Fig. S6). In addition to the fossil-based calibrations adapted from published studies, we used the posterior age distributions derived from the first-step dating analysis to calibrate the overlapping nodes in the species tree containing both eukaryotes and Cyanobacteria (Fig. S1c; Table S1; Table S2). Note that we placed the cyanobacterial-fossil based calibrations at the total group of Nostocales and Pleurocapsales by taking into account the incomplete taxon sampling of their early-split lineages, as well as the possibility that their fossils belong to the stem lineages rather than the crown groups. In this way, we allow for the posterior ages of crown Nostocales and Pleurocapsales groups to be younger than the known fossil records.

Our molecular dating analysis was conducted using MCMCTree with the best-fit clock model, which was selected using the program “mcmc3r” following published studies (McGowen et al., 2019; Wang and Luo, 2021). For calibrations and species tree topology that remain disputed, we took all the possibilities in our analysis for comparison (Fig. S4). We also tested the necessity for using Bayesian sequential dating approach and hard bound calibration strategy in our analysis (Fig. S7; see details in Supplementary Discussion) and tested the convergence of the posterior age estimates (Fig. S8).

The gains and losses of *Prochlorococcus* orthologous gene families were inferred with a gene tree-species tree reconciliation approach according to our recent studies (Zhang et al., 2024). The best-fit reconciliation tool was selected through a comprehensive simulation-based benchmarking analysis [see Fig. S5 in (Zhang et al., 2024)]. In our implementations, we performed the reconstruction first without using the SAGs to avoid the false prediction of genomic events due to the relatively low completeness of the SAGs (e.g., 81.9% for AMZI-B-ETNP) and then used the same reconstruction methods to validate these evolutionary events when SAGs were included.

The reduction of *Prochlorococcus* selection efficiency in ancient time was inferred by comparing the genome-wide *d*_R_/*d*_C_ (i.e., the relative rate of radical versus conservative nonsynonymous substitutions) between the target lineage and its sister lineage based on our recent study (Luo et al., 2017) (see details in Supplementary Methods).

## Code availability

The custom scripts used to analyze the data are available in the online GitHub repository (https://github.com/luolab-cuhk/Prochl-NOE).

## Acknowledgements

This work is supported by the Natural Science Foundation of China (42293294), the Hong Kong Research Grants Council (RGC) General Research Fund (GRF) (14110820), the Natural Science Foundation of Guangdong Province, China (2022A1515010844 to HZ), the China Postdoctoral Science Foundation (2021M702296 to HZ), and the CUHK Direct Grant (4053551).

## Notes

https://github.com/luolab-cuhk/Prochl-NOE

### Competing Interest Statement

The authors have declared no competing interest.

### Summary of Updates

The structure of the main text has been revised. Supplementary file updated.

