## Supplementary material for "Emergence of *Prochlorococcus* in the Tonian oceans and the initiation of Neoproterozoic oxygenation": SI

**This PDF file includes:**

Supplementary Results
Supplementary Discussion
Supplementary Methods
References
Fig. S1 to S8
Table S1 to S2

### Supplementary Results

<u>Using alternative calibration and alternative species tree topology in dating analysis</u>

To test the robustness of the evolutionary timeline of *Prochlorococcus* estimated with the focal strategy, we evaluated the impact of different fossil records on *Prochlorococcus* divergence time estimates. By employing alternative lower bound calibrations of the total Nostocales group (1.2 Gya and 2.0 Gya instead of 1.6 Gya) and alternative lower bound calibrations of the crown Rhodophyta group (1.047 Gya and 1.6 Gya instead of 1.2 Gya) (see calibration information in Supplementary Methods), we found that the posterior ages of Proch-AMZI/II/III-LCA, Proch-AMZI/II-LCA and SBE-LCA were largely consistent with that estimated with the focal strategy (Fig. S3). Note that the posterior ages of the crown Nostocales group were younger than the fossil records. The discrepancy is not unexpected since the fossil calibration is placed on the total group, as also found in the timetree of animals shown in the study (dos Reis et al., 2015) (e.g., Tetrapoda, Osteichthyes, Amniotes, etc.). This suggests either incomplete taxon sampling of early-split lineages of Nostocales or that the fossils (Horodyski and Donaldson, 1980; Amard and Bertrand-Sarfati, 1997; Tomitani et al., 2006) are derived from the stem lineage rather than the crown group of Nostocales.

We also tested the effect of using alternative tree topological structures on posterior age estimates. In the eukaryotic portion of the tree, Glaucophyta was thought to represent the earliest-split branch of the photosynthetic eukaryotes in some phylogenomic studies (Topo-1 in Fig. S1a) (Parfrey et al., 2011; Burki et al., 2012; Ponce-Toledo et al., 2019a), but Rhodophyta takes the basal position in others (Topo-2 in Fig. S1a) (Schön et al., 2021). We thus used an alternative eukaryotic species tree topology for molecular clock analyses, in which Rhodophyta, instead of Glaucophyta, was positioned at the base of the eukaryotic portion of the species tree

(Strassert et al., 2021a). Likewise, we used an alternative eukaryotic species tree topology in which the Prasinodermophyta group locates at the base of Chlorophyta instead of the base of Viridiplantae, as that indicated by the recent study (Strassert et al., 2021b) (Topo-3 in Fig. S1a). We found that the posterior ages estimated with these alternative settings are largely consistent with that estimated with the focal strategy.

Using different clock models in dating analysis

While our clock model selection is more supportive of an independent rate (IR) model for the 12-gene dataset used in the focal strategy ("T30"; see details in Supplementary Method), we note that the auto-correlated rate (AR) model cannot be fully rejected (7 versus 5 gene families support the IR and AR model, respectively; Supplementary Data 2). By applying the AR model, we found that the posterior ages of the Proch-AMZI/II/III-LCA (978 Mya; 95% HPD: 1,117-858 Mya) and Proch-AMZI/II-LCA (810 Mya; 95% HPD: 938-690 Mya) were largely consistent with that estimated with the IR model. The SBE-LCA (466 Mya; 95% HPD: 572-356 Mya), became younger than that estimated with the IR model, but the branch leading to the SBE-LCA still encompasses the Snowball Earth events (Fig. S3; Fig. S4).

Using different molecular datasets in dating analysis

To reduce the bias in posterior age estimates introduced by using a limited number of genes, we compiled a larger dataset comprising 19 orthologous gene families ("T40"; Supplementary Data 2). While the AR model gained more support in this case, the IR model cannot be fully rejected (8 versus 11 gene families supporting the IR and AR model, respectively; Supplementary Data 2). We found this 19-gene set ("T40") gave very different posterior age estimates than the 12-gene set ("T30"). When AR was used, for example, age estimates of Proch-AMZI/II/III-LCA, Proch-AMZI/II-LCA and SBE-LCA remarkably shifted toward the past by using the 19-gene set ("T40") and were no longer overlapped with those estimated with the 12-gene set ("T30"). When IR was used instead, the posterior mean age shifted toward the past by using the 19-gene set ("T40") to a lesser extent, and the uncertainties of the posterior ages (indicated the by the 95% HPD interval) were enlarged (Fig. S3). These problems appear to be associated with the use of two problematic genes, as a few *Prochlorococcus* SAGs were clustered with the anoxygenic Vampirovibrionia and Sericytochromatia rather than the remaining *Prochlorococcus* (Fig. S6). By leaving these two genes out ("T40*"), both AR and IR models gave posterior ages more consistent with those based on the "T30" dataset (Fig. S3). Further removal of another three gene families in which *Prochlorococcus* does not form a monophyletic group (Fig. S6) led to the dataset "T40**", based on which the posterior age estimates remain consistent with that estimated with the dataset "T30" (Fig. S3).

<u>Dating the divergence time of *Prochlorococcus* without the use of Bayesian sequential approach and hard bound calibration</u>

In addition to the more extensive taxon sampling including the eukaryotic lineages that have undergone secondary endosymbiosis, our plastid-based dating strategy differs from the previous ones (Shih et al., 2016; Sánchez-Baracaldo et al., 2017; Fournier et al., 2021) in two aspects. First, we implemented the Bayesian sequential dating approach to provide a more accurate time estimate for eukaryotic lineages. Without this improvement, the posterior ages of the crown eukaryotic group (2.38 Gya; 95% HPD: 2.21-2.64 Gya) would have been much more ancient than published estimates (appr. 1.87-1.68 Gya) (Fig. S7a) (Parfrey et al., 2011; Betts et al., 2018;

Wang and Luo, 2021). Second, we implemented hard bound calibrations to avoid the potential bound violation problem. If the soft bound calibrations were used, the posterior ages of total Nostocales group (0.99 Gya; 95% HPD: 1.166-0.806 Gya) and total Pleurocapsales group (1.14 Gya; 95% HPD: 1.238-1.031 Gya) are far younger than the limits of their minimum age constraints (1.6 Gya and 1.7 Gya, respectively) (Fig. S7b).

### Supplementary Discussion

<u>Different *Prochlorococcus* divergence time in the present and the previous studies</u>

Previous molecular clock studies, although not focusing on *Prochlorococcus*, have estimated the evolutionary timeline of Cyanobacteria. Given the variable taxon sampling of the *Prochlorococcus* lineages in those studies, we compared the posterior ages of the total *Prochlorococcus* group among these studies and divided them into the older and the younger-age groups. The former includes the age estimates at ~820 Mya (Sánchez-Baracaldo et al., 2014), ~840 Mya (for crown *Prochlorococcus* group due to the unavailability of the age for total *Prochlorococcus* group in this study) (Sánchez-Baracaldo, 2015) and ~900 Mya (Schirrmeister et al., 2015; Zhang et al., 2021), which are largely consistent with that estimated in the present study (959 Mya; 95% HPD: 846-1,077 Mya; Fig. 1). The latter group includes the age estimates at ~200 Mya (Boden et al., 2021; Martinez-Gutierrez et al., 2023), ~400 Mya (Schirrmeister et al., 2013; Sánchez-Baracaldo et al., 2019; Fournier et al., 2021) and ~500 Mya (Sánchez-Baracaldo, 2015; Sánchez-Baracaldo et al., 2017). By looking into these younger-age group studies, we found that most of them using SSU and LSU rRNA genes in their dating analysis (Schirrmeister et al., 2013; Sánchez-Baracaldo, 2015; Sánchez-Baracaldo et al., 2019; Boden et al., 2021). Fournier et al. (2021) and Martinez-Gutierrez et al. (2023) recently used an enlarged ribosomal protein dataset in their dating analysis but also estimated early ages of the total *Prochlorococcus* group, suggesting that other factors might have contributed to the large discrepancies in posterior ages between the present study and the younger-age group studies.

In Martinez-Gutierrez et al. (2023), the calibration settings may be inappropriate. First of all, they constrained the origin of aerobic Nitrososphaerales, aerobic Marinimicrobia, and nitrite oxidizing bacteria groups after the GOE without the consideration of the known oxygen production before the GOE (Crowe et al., 2013; Planavsky et al., 2014; Lalonde and Konhauser, 2015). Next, the rationale for using the age of the most ancient record of methane as the minimum time constraint for Bacteria and Archaea is unclear, considering that the possibility of an abiotic source of methane on early Earth cannot be ruled out (Ueno et al., 2006; Alleon and Summons, 2019). Finally, the commonly used cyanobacterial fossils were not adopted in that study, which likely biased the posterior age estimation of cyanobacterial groups.

By taking Fournier's study as another example, we compared the methodological differences between the two studies in detail and found that the major difference comes from the use of clock model. Fournier et al. adopted the three clock models implemented in the software PhyloBayes, including the UGAM model, the LN model and the CIR process model. The UGAM model parallels the IR model implemented in MCMCTree, and the main difference between them is that the former uses a Gamma distribution while the latter uses a log-normal distribution to model the rate variation across branches. The LN model is an auto-correlated log-normal model (or geometric Brownian model), which is equivalent to the AR model implemented in MCMCTree. The CIR process model is also an auto-correlated model but uses stationary distribution instead of the lognormal distribution (Lepage et al., 2007). This model has not been implemented in MCMCTree. In contrast to the present study, in which the use of AR clock model (1,110 Mya; 95% HPD: 982-1,248 Mya) resulted in slightly more ancient age estimates for the total *Prochlorococcus* group than the use of IR clock model (960 Mya; 95% HPD: 846-1,077 Mya), we found that in Fournier's study, the use of UGAM clock model (736 Mya; 95% HPD: 425-1,169 Mya) resulted in much more ancient age estimates than the use of LN model (316 Mya; 95% HPD: 211-479 Mya) and CIR process model (417 Mya; 95% HPD: 326-575 Mya). Therefore, an unbiased evaluation of the difference clock model is required in their study.

It is well known that model selection for Bayesian model clock analysis should be performed by Bayesian model selection (Lartillot et al., 2009; dos Reis et al., 2018). In our MCMCTree analysis, we used the stepping-stone method implemented in the program "mcmc3r" to perform this analysis. Similar statistical tools are available in PhyloBayes, which rely on the calculation of Bayes factor under the normal approximation, cross-validation and posterior predictive testing (Lartillot et al., 2009). Unexpectedly, Fournier et al. did not perform this well-established statistical method. Instead, they chose the clock model that is most compatible with the identified horizontal gene transfer (HGT) events according to the idea that the donor species must be older than the recipient species. Based on this rationale, the CIR process model was chosen because the posterior age estimates resulting from this model were most compatible with identified HGT events (75.79%), whereas posterior ages resulting from using the UGAM and LN models were less compatible with the HGT events (51.83%-58.85%) (Fournier et al., 2021).

There are several important problems with the way Fournier et al. used to select clock model. First, as we interpret it, the idea of using HGT-based time constraints is to leverage the relative divergence order of the gene donor and the gene recipient to accept or reject the posterior samples in an MCMC run. Accordingly, it is "expected" that there are some conflicts between the relative divergence order implied by the identified HGT events and the posterior ages estimated without considering HGT, so that the HGT-based relative time constraints can help to "correct" the posterior age estimation (Davín et al., 2018; Magnabosco et al., 2018). In this way, however, the clock model with the highest compatibility is least "useful" as it did not contribute much to correcting the posterior age estimation. Hence, using the compatibility as support for model selection appears to be somehow self-contradictory. Next, the way Fournier et al. used for clock model selection does not contain a statistical component. There was no test to support that the CIR model with 75.79% compatibility was significantly better than the UGAM and LN model with 51.83%-58.85% compatibility.

In addition to the clock model, there are other parameter differences between the present and Fournier's study. In terms of the calibration, for example, we adopted a few more calibrations on the important eukaryotic groups, including the Chlorophyta group, the Embryophyta group and the Spermatophyta group, which were omitted in Fournier's study. Moreover, we implemented Bayesian sequential dating approach to provide more informative time prior to the crown eukaryotic group, which in Fournier' s study was constrained by a secondary calibration (i.e., <1.8 Gya for crown eukaryotic group according to previous studies). In terms of the molecular dataset, we manually screened the orthologous gene families based on their verticality (indicated by $\Delta$LL, i.e., the likelihood difference of the gene tree reconstructed with and without the constraint of the species tree topology; see Supplementary Methods for details) and the evolutionary rate difference (indicated by the branch length difference between the cyanobacterial group and the eukaryotic group; see Supplementary Methods for details), while Fournier et al. directly used the 30 conservative large and small subunit ribosomal proteins (originally used for species tree reconstruction) without further examination. While these methodological differences seem to be critical for molecular dating analysis, they indeed have less influence on posterior age estimates compared to the choice of clock model. Regardless of these methodological differences, we found that the posterior ages of the total *Prochlorococcus* group estimated under the UGAM model in Fournier's study (736 Mya; 95% HPD: 425-1,169 Mya) overlapped with that estimated in the present study (960 Mya; 95% HPD: 846-1,077 Mya under the IR model), while the 95% HPD interval of the former is very large.

<u>Comparing the divergence time of the crown group Cyanobacteria between the present and previous studies</u>

The origin of Cyanobacteria has been a topic of extensive discussion and is generally accepted to have occurred at ~2.4 Gya during the Great Oxidation Event (GOE) (Schirrmeister et al., 2016). Using 2-methylhopane as a biomarker for Cyanobacteria, their origin has been traced back to approximately 2.7 Gya (Brocks et al., 2003). However, this connection between 2-methylhopane and Cyanobacteria faced challenges when it was discovered that 2-methylhopane can be produced by the anoxygenic phototroph *Rhodopseudomonas palustris* under anaerobic conditions (Rashby et al., 2007). A more recent study based on Mo isotope data proposes that oxygenic photosynthesis played an important role in oxygen accumulation in shallow marine environments hundreds of millions of years before the GOE, dating back to ~2.95 Gya (Planavsky et al., 2014). Likewise, an independent study utilizing U-Th-Pb isotope data suggests the existence of a distinct redoxcline between deep and shallow seawater at approximately 3.2 Gya (Satkoski et al., 2015). The enrichment of oxygen in the upper water column has been attributed to the activities of oxygen-producing microorganisms, suggesting that Cyanobacteria have evolved prior to 3.2 Gya (Satkoski et al., 2015).

While geological evidence strongly supports an early origin of Cyanobacteria, many recent molecular clock analyses have yielded younger age estimates for the crown Cyanobacteria group (Sánchez-Baracaldo et al., 2022). For example, Sánchez et al. estimated the posterior mean age of the crown Cyanobacteria group at ~2.6 Gya and ~2.9 Gya when the upper bound limit of the crown Cyanobacteria group was set to 2.7 Gya and 3.0 Gya, respectively (Sánchez-Baracaldo, 2015). Sánchez et al. then expanded the taxonomic sampling by including Archaeplastida lineages and the non-oxygenic cyanobacterial lineages and integrating additional fossil-based calibrations, such as eukaryotic fossil-based calibrations and endosymbiont-based calibrations, in their most recent studies (Sánchez-Baracaldo et al., 2017; Boden et al., 2021). These studies, however, arrived at similar estimates for the divergence time of crown Cyanobacteria group (i.e., ~2.6 Gya and ~2.9 Gya when the upper bound limit of crown Cyanobacteria group was set to 2.7 Gya and 3.0 Gya, respectively). These priors impose a strong limit for the time of the crown Cyanobacteria group, but there is not a very good reason to rule out the possibility of their earlier origin.

We noticed that the posterior mean ages of the crown Cyanobacteria group in abovementioned studies are very close to their upper bound limits, suggesting that the posterior age of the crown Cyanobacteria group was strongly affected by the prior particularly the upper time bound, which however was often set in an arbitrary way. Hence, a more relaxed thus unbiased time prior for Cyanobacteria dating analysis is required. Schirrmeister et al. elevated the upper bound limit of the crown Cyanobacteria group to 3.85 Gya, which assumes that Cyanobacteria could have originated after the Late Heavy Bombardment at 3.85 Gya (Schirrmeister et al., 2015). In this way, the crown Cyanobacteria group was dated back to 3.3 Gya and 3.6 Gya based on different inner node calibrations (Schirrmeister et al., 2015). To relax the time prior for crown Cyanobacteria group, an alternative calibration strategy has been proposed, which imposes the time constraint at 3.0 Gya (the time when free oxygen was available in water column) as the lower bound age for the total Cyanobacteria group, rather than as the upper bound age for the crown Cyanobacterial group (Zhang et al., 2021). In this way, the posterior mean age of the crown Cyanobacteria was estimated to ~3.4 Gya (Zhang et al., 2021).

To avoid using the calibrations on specific cyanobacterial lineages to estimate their own divergence time, Garcia et al. chose not to impose any calibrations on the total or crown Cyanobacteria group. Instead, they relied on the internal node calibrations on filamentous Cyanobacteria and heterocyst-forming Cyanobacteria for their dating analysis (Garcia-Pichel et al., 2019). When the root maximum age was set to 4.5 Gya, they found the posterior mean age of crown Cyanobacteria group at ~3.6 Gya (Garcia-Pichel et al., 2019). Likewise, Fournier et al. employed the internal node calibrations on microbial mat-forming Cyanobacteria, filamentous Cyanobacteria and endolithic Cyanobacteria in their dating analysis and avoided using the time constraints on either the total or crown Cyanobacteria group. In contrast to Garcia et al., 2019, Fournier et al. incorporated additional bacterial calibrations to their dating analyses and then estimated the posterior mean age of the crown Cyanobacteria group at 3.1-3.6 Gya under different clock model (Fournier et al., 2021). When eukaryotic fossil-based calibrations were further added, the posterior mean ages of the crown Cyanobacteria group were 2.9-3.2 Gya under different clock models (Fournier et al., 2021). The above illustrates the use of an uninformative time priors could lead to less biased time estimates for dating deep phylogenies, as also observed in other studies (Wang et al., 2020).

Collectively, it appears that the younger age estimates for the crown Cyanobacteria group may have been caused by using an inappropriate upper bound limit. When more relaxed calibrations were applied to the crown Cyanobacteria group, their divergence time became considerably older. In the present study, we did not impose any maximum age constraint on the crown or total Cyanobacteria group. Instead, we constrained the lower bound limit of the total Cyanobacteria group using the time of GOE while leaving its upper bound limit open. In this way, when the root maximum age changed from 3.8 Gya to 4.5 Gya, the posterior ages of the crown Cyanobacteria group changed from 3.19 Gya (95% HPD: 2.95-3.42 Gya) to 3.52 Gya (95% HPD: 3.87-3.18 Gya) (Fig. S5a), which are very close to that estimated in the published studies with elevated or unlimited upper bound limit of the crown Cyanobacteria (Schirrmeister et al., 2015; Garcia-Pichel et al., 2019; Zhang et al., 2021). For test, we further increased the root maximum age from 4.5 Gya to 5.5 Gya, although these ages have no biological or geological significance. By doing this, we found that the posterior ages of the crown Cyanobacteria group became even older (3.95 Gya; 95% HPD: 3.40-4.52 Gya) (Fig. S5a). Intriguingly, in these tests, the origin time of *Prochlorococcus* (Prochl-AMZI/II/III-LCA) remained largely consistent at ~900 Mya, regardless of whether the root maximum age was set to 3.8 Gya or 5.5 Gya (Fig. S5a). It appears that the cyanobacterial lineages that subtend the plastid lineage can be well constrained using the plastid-based molecular clock, whereas those that branched off earlier than the plastid lineage cannot. This hypothesis is supported by Fig. S5b, where the trend line deviates from the y=x line at ~1.8 Gya, close to the time when the crown eukaryotic group evolved.

### Supplementary Methods

A) Plastid-based Bayesian sequential dating strategy

A1) Taxon sampling and data collection

To initiate the plastid-based molecular dating strategy, we obtained plastid-encoded protein sequences of 34 representatives of eukaryotic algae (Chlorophyta, Charophyta, Cryptophyta, Diatom, Dinoflagellate, Rhodophyta, Glaucophyta) and land plants (Embryophyta) from the NCBI Refseq release of plastid database (March 2022) (https://ftp.ncbi.nlm.nih.gov/refseq/release/plastid/) (Supplementary Data 1). Among the 34 representative eukaryotes, genomic data are available for 15 species in Dicots PLAZA (5.0) database (https://bioinformatics.psb.ugent.be/plaza/) (Supplementary Data 1). Four phytoplankton species, including *Emiliania huxleyi*, *Phaeodactylum tricornutum*, *Thalassiosira pseudonana*, and *Aureococcus anophagefferens*, were excluded from them in our dating analysis due to the incongruence of their phylogenetic positions inferred based on nuclear-encoded and plastid-encoded proteins (Morden and Sherwood, 2002; Ponce-Toledo et al., 2019b), as their plastids are acquired by a secondary or even tertiary endosymbiosis events (Strassert et al., 2021b).

We also obtained the genomic sequences of the 159 oxygenic Cyanobacteria and the 8 anoxygenic Vampirovibrionia (formerly known as Melainabacteria) and Sericytochromatia from GenBank by following our recent publication (Zhang et al., 2021) (Supplementary Data 1). Previous studies identified the uncultivated *Prochlorococcus* clades LLV and LLVI based on the 16S-23S rRNA internal transcribed spacer (ITS) sequences (Lavin et al., 2010). Since both clades inhabit the nutrient-rich and oxygen-depleted layer of anoxic marine zone (AMZ), they were recently renamed as AMZI and AMZII, respectively (Ulloa et al., 2021). Single-cell amplified genomes (SAGs) are now available to both AMZI and AMZII clades, as well as a novel clade AMZIII, which locates at the base of the phylogenomic tree of *Prochlorococcus*, representing the closest relatives to *Synechococcus* (Ulloa et al., 2021). We adopted five

*Prochlorococcus* SAGs in our dating analysis, including AMZI-A-ETSP, AMZI-B-ETNP, AMZI-B-ETSP, AMZII and AMZIII (Ulloa et al., 2021) because their completeness was at least 80% (Supplementary Data 1) according to CheckM v1.0.13 (Parks et al., 2015).

A2) The first-step Bayesian sequential dating analysis

Bayesian sequential dating approach was developed a decade ago (dos Reis et al., 2012); it did not gain much attention until a recent study where it was employed in estimating a species-level evolutionary timeline of mammals (Álvarez-Carretero et al., 2022). By implementing a small-scale (first-step) dating analysis (with a few taxa but many genes) followed by a full-scale (second-step) dating analysis (with numerous taxa but a reduced number of genes not overlapped with the genes used in the first step), this approach is able to reduce the computational cost and increase the accuracy for dating a large dataset with numerous taxa (dos Reis et al., 2012). The key step in the Bayesian sequential method is using posterior age distributions in the small-scale dating analysis as time priors at overlapping nodes in the full-scale dating analysis (dos Reis et al., 2012).

Here, we implemented the Bayesian sequential dating analysis to provide a more accurate time estimate for photosynthetic eukaryotic lineages in Cyanobacteria dating analysis. The molecular data used in the first step of sequential dating analysis is composed of nuclear-encoded proteins of the 11 eukaryotes with genomic data.

We identified 251 orthologous gene families shared by the 11 eukaryotes by searching against a pre-identified eukaryote-wide dataset that contains 320 manually curated marker nuclear genes (Strassert et al., 2021b). The mean average evolutionary rate of each gene family was estimated with CODEML in PAML package under the LG + Γ4 + F model. The 251 gene families were then partitioned into 1, 3, 5, 7 and 9 partitions based on the gene evolutionary rate using Gaussian Mixture Model (GMM) clustering method implemented in the R package ClusterR (Mouselimis, 2023). Partitioned datasets were then passed to MCMCTree together with the eukaryotic subtree for posterior age estimation. In general, the posterior ages of eukaryotic lineages were consistent, except for the case where a single partition and a single substitution model was used, which did not fit the data well as indicated by its larger Akaike Information Criterion (AIC) value (Fig. S1b).

Next, we identified the best-fit posterior age distribution from all the available distributions implemented by MCMCTree, namely skew-normal, skew-t and Gamma distributions, for each ancestral node based on Akaike Information Criterion (AIC), which was similarly done in a recent study to date mammal evolution (Álvarez-Carretero et al., 2022). Among these three distributions, the skew-normal and skew-t distributions were used to fit the outlying observations showing departure from the normal distribution and the student's t-distribution, respectively (Azzalini and Genton, 2008). The approximated distributions were then mapped back to the corresponding nodes of the tree containing both Cyanobacteria and plastids as the time prior for the second-step Bayesian dating analysis (Fig. S1; Table S2). To assess the effectiveness of Bayesian sequential approach in the present study, we compared the posterior ages of the eukaryotic nodes obtained in the first-step sequential dating analysis to the effective time prior ("usedata = 0" in the MCMCTree configuration file) in corresponding nodes in the second-step sequential dating analysis. We found that when the 5-, 7-, and 9-partition data were used, the distributions of posterior age and the corresponding effective prior were highly consistent, suggesting that the Bayesian sequential dating method works well for these analyses (Fig. S2).

A3) The second-step Bayesian sequential dating analysis

The small-scale and the full-scale analyses in the sequential dating process must use different molecular data, otherwise the duplicate use of overlapped genes would lead to a squaring of the likelihood and consequently converge to a wrong posterior age estimates (Rannala and Yang, 2007; dos Reis et al., 2018). Therefore, the molecular data used in the second step of sequential dating analysis was composed of plastid-encoded proteins of the 34 eukaryotes and their orthologues in the five *Prochlorococcus* SAGs and the 167 cyanobacterial isolates. These orthologous gene families were identified by searching against a dataset containing 97 plastid marker genes using BLASTP with an e-value of 1e-30 (Supplementary Data 2) (Ponce-Toledo et al., 2017). We chose this dataset for two reasons. First, it was compiled based on a broad and balanced taxonomic sampling of Cyanobacteria and Archaeplastida (a major eukaryotic group comprising photoautotrophic red algae, green algae, land plants, and glaucophyta). Second, the phylogenetic tree of plastid marker genes were manually inspected to remove those showing evidence of horizontal gene transfer (HGT) among cyanobacterial species (Ponce-Toledo et al., 2017).

A3.1) Gene dataset compilation for molecular clock analysis

Since the genes used for molecular clock analysis may affect the posterior age estimates, we screened the orthologous gene families by considering two major factors. First, genes that show important phylogenetic incongruence with the species tree due to HGT or other evolutionary processes are not appropriate for molecular dating. HGT events are known to change the branch lengths of the nodes involved in HGT while leaving other branches largely unaltered in their lengths (Moody et al., 2022). Such local branch length changes are expected to affect the age estimation, as opposed to the potentially limited impact imposed by global branch length changes that are often caused by using inadequate amino acid replacement models (Tao et al., 2020). Second, genes that show very large evolutionary rate variation are also not appropriate. We note that MCMCTree models the evolutionary rate difference across lineages (also known as evolutionary rate drift) by a log-normal distribution where the default values of the parameter $\sigma^2$ which governs the extent of across-lineage rate heterogeneity (sigma2_gamma in mcmctree) has been shown to already accommodate large rate variation (Brown and Yang, 2011). In our dataset, however, plastid-encoded genes in photosynthetic eukaryotes appear to have much higher rates than their orthologous genes in Cyanobacteria (Ponce-Toledo et al., 2017), which makes it possible that the rate variation modeled by MCMCTree is not adequate to account for the rate variation contained in our dataset.

We developed a bioinformatics procedure to deal with these problems. In brief, we aligned the amino acid sequences of the 97 orthologues with MAFFT v7.453 under the auto-determined mode (Katoh et al., 2002) and trimmed the alignments using trimAl v1.4 with the parameter "automated1" (Capella-Gutiérrez et al., 2009). For each orthologous gene family, we constructed the maximum likelihood tree using IQ-TREE v2.0.6 (Minh et al., 2020) under the discrete Gamma rate model and the auto-determined amino acid substitution model either with or without the backbone species tree topology. Note that the backbone species tree used here was adapted from the general consensus phylogeny of eukaryotes and Cyanobacteria in published studies (Ponce-Toledo et al., 2017; Cheng et al., 2019; Gulbrandsen et al., 2021; Strassert et al., 2021b; Su et al., 2021; Ulloa et al., 2021; Zhang et al., 2021) and was pruned to sample the same taxa as each gene tree. By comparing the log-likelihood values of the gene tree constructed with and without the backbone species tree topology, we ranked the 97 orthologous gene families by ΔLL (a proxy for the extent to which a marker gene rejects vertical evolution) (Moody et al., 2022) (Supplementary Data 2). By comparing the root-to-tip distance among species (Moody et al., 2022), we ranked the 97 orthologous gene families by evolutionary rate difference between Cyanobacteria and eukaryotes (Supplementary Data 2). We then compiled two datasets comprising 12 and 19 orthologous gene families that ranked in the top (best) 30% and 40%, respectively, according to both ΔLL and evolutionary rate difference. The former was marked as "T30" and was used as the focal dataset in our analysis. The latter was marked as "T40" and was used as an alternative dataset for tests.

A3.2) Molecular clock model selection

MCMCTree implements two relaxed clock models for divergence time estimation, the independent-rates (IR) model and the autocorrelated-rates (AR) model. The marginal likelihood value of the MCMC analysis can be used to select the best-fit clock model, which is calculated based on the exact likelihood values sampled during the MCMC analysis. However, this calculation is computationally expensive, especially for large datasets (McGowen et al., 2019). As the size of the dataset used for plastid-based dating analysis was increased by incorporating eukaryotic species, an unbiased smaller dataset was compiled for clock model selection; this strategy follows the published studies (McGowen et al., 2019; Wang and Luo, 2021). Specifically, we selected 18 cyanobacterial representatives by clustering the leaves with the software TreeCluster based on phylogenetic distance (Balaban et al., 2019) ("max clade" method; branch length threshold = 1; Supplementary Data 1) in combination with the 11 eukaryotic species with available genomic DNA sequences for clock model test (Supplementary Data 1). As the exact likelihood value of the MCMC analysis can only be calculated from nucleotides, we recoded the amino acids in our molecular data into four nucleotide characters according to their physicochemical properties with the Dayhoff4 recoding scheme by using an in-house script (Wang and Luo, 2023). We then employed the R package "mcmc3r" to estimate the Bayes factor and the Posterior Probability of the IR and AR model for each gene family using the stepping-stones method (Supplementary Data 2) (dos Reis et al., 2018; Álvarez-Carretero et al., 2022).

A4) Calibration information

Non-uniform prior distribution, like truncated Cauchy distribution, was used to describe the calibration density relative to the minimum constraint alone and was known to be sensitive to parameter choice (Warnock et al., 2012). By contrast, uniform priors allow the user to accommodate a view that nothing is known about the divergence time relative to the constraints and thus reduced the artificial bias in the prior of calibration (Warnock et al., 2015). Following published studies (Parfrey et al., 2011; Betts et al., 2018; Wang and Luo, 2021), we provided an Earth age-based upper bound limit (4.5 Gya) to those calibrations without fossil-based maximum age constraint. Calibrations with both lower and upper bounds in MCMCTree follow a uniform-like distribution [B(*tL*, *tU*, *pL*, *pU*)], in which the parameters *tU*, *tL*, *pU*, and *pL* represent the upper bound age, the lower bound age, the right tail probability (i.e., the probability of upper bound violation) and the left tail probability (i.e., the probability of lower bound violation), respectively (Yang and Rannala, 2005). Since correctly identified fossils are known to provide the confirmation that a lineage existed at a certain time without any ambiguity (Benton et al., 2007; Donoghue and Benton, 2007), we constrained all the fossil-based minimum age constraints and the Earth age-based maximum age constraint (4.5 Gya) as hard bound (*pL*=1e-300, *pU*=1e-

300) to prevent the bound violation problem while keeping the fossil-based maximum constraints as soft bound. Note that the node numbers correspond to those in Fig. S1a or Fig. S1c.

Node_1: Root

Minimum age: 2.32 Gya

Maximum age: 4.5 Gya

Justification: MCMCTree requires a root maximum age for dating analysis (Yang and Rannala, 2005). To avoid the bias artificially induced by constraining the age prior, we set the root maximum age to 4.5 Gya when the planet Earth formed (Allegre et al., 1995). The minimum age of the root was set to the age of Great Oxidation Event (GOE) at 2.32 Gya (Bekker et al., 2004), which is a conservative setting and was used as the minimum age constraint for the total oxygenic Cyanobacteria (see details in Node_2).

Node_2: Total oxygenic Cyanobacteria

Minimum age: 2.32 Gya

Maximum age: 4.5 Gya

Justification: The minimum age of the oxygenic Cyanobacteria was inferred from the GOE, which is considered as the result of Cyanobacterial oxygenic photosynthesis (Kopp et al., 2005; Schirrmeister et al., 2013). GOE has been used to constrain the minimum age of the crown oxygenic Cyanobacteria group in published studies (Sánchez-Baracaldo et al., 2014; Sánchez-Baracaldo et al., 2017; Wolfe and Fournier, 2018). However, the identification of non-oxygenic Cyanobacteria suggests that oxygenic photosynthesis may have evolved at the stem linage of oxygenic Cyanobacteria. Consequently, by following our recent study (Zhang et al., 2021), we constrained the lower bound of the total oxygenic Cyanobacteria group at 2.32 Gya. The maximum age of the total oxygenic Cyanobacteria was set to 4.5 Gya due to the absence of paleontological or paleogeologic evidence.

Node_3: Total group of Pleurocapsales

Minimum age: 1.7 Gya

Maximum age: 4.5 Gya

Justification: The minimum age of total Pleurocapsales group was constrained at 1.7 Gya based on the microfossil of Pleurocapsales found in Hebei, China (Zhang and Golubic, 1987). We note that the same fossil has been used to constrain the minimum age of Pleurocapsales on their crown group (Sánchez-Baracaldo et al., 2017). However, given the fact that the apomorphic characters must have evolved earlier than the divergence of the crown group of assigned lineage (Marshall, 2019), a more secure way to apply these morphological fossils is to constrain the total group instead of the crown group of Pleurocapsales. The same calibration strategy has been implemented in published studies (Wang and Luo, 2021; Zhang et al., 2021).

Node_4: Total Nostocales group

Minimum age: 1.2 Gya, 1.6 Gya and 2.0 Gya

Maximum age: 4.5 Gya

Justification: Akinete is known as the feature of Nostocales, which allows for the survival of the cell under extreme environmental conditions (Tomitani et al., 2006). The minimum age of the Nostocales group was set to 1.6 Gya based on the akinete fossil preserved in McArthur Group of Northern Australia (Tomitani et al., 2006). The alternative minimum age constraint of the

Nostocales group was inferred based on the fossil of coccoid cyanophyte in the 1.2 Gya Dismal Lakes Group of Northwest Canada (Horodyski and Donaldson, 1980), and the akinete fossil identified in the stromatolites in the 2.0 Gya Franceville Group, Gabon (Amard and Bertrand-Sarfati, 1997). Both these alternative minimum age constraints have been used in the published study (Wolfe and Fournier, 2018). For the same reason mentioned above (Node_2), we constained the miminum age of Nostocales on their total group.

Node_5: Crown Viridiplantae group & Node_6: Crown Chlorophyta group

Minimum age: 0.947 Gya

Maximum age: 1.891 Gya

Justification: *Proterocladus antiquus* represents the earliest green seaweed to our best knowledge. The recently discovered microfossil from the Nanfen Formation in North China was suggested as the benthic chlorophyte *P. antiquus* based on the morphological characters for example, the siphonocladous construction and the branching and holdfast structures of the cell (Tang et al., 2020). While the age of the Nanfen Formation has not been directly inferred from the isotope data, the age of the overlying Qiaotou Formation was dated at ~0.947 Gya (Zhao et al. 2020). We thus constrained the minimum age of the crown Viridiplantae/ Chlorophyta group conservatively at 0.947 Gya.

The well-preserved acritarchs with large size and fine-scale morphological complexity in the shales of Changzhougou Formation in China (1.823±0.068 Gya) were considered as the simple eukaryote that evolved earlier that all other reported multicellular eukaryotes (Lamb et al., 2009). By following the published study (Morris et al., 2018), we constrained the maximum age of the crown Viridiplantae/Chlorophyta group at 1.891 Gya.

Node_7: Crown Embryophyta group

Minimum age : 0.45 Gya

Maximum age: 0.509 Gya

Justification: Trilete spores were considered as the feature restricted to crown Embryophyta and thus their fossils were used to constrain the minimum age of the crown Embryophate group in published studies (Clark and Donoghue, 2017; Wang and Luo, 2021). Consequently, we constrained the minimum age of the crown Embryophyta group by using the oldest fossil of the trilete spores identified in the Quasim Formation of northern Saudi Arabia (448.5Mya) (Gradstein et al., 2012). The same calibration strategy has been implemented in published studies (Betts et al., 2018).

We constrained the maximum age of the crown Embryophyta group at 509 Mya based on the age of Bright Angel Shale in the eastern Grand Canyon, Arizona, USA (507.2-509 Ma) (Baldwin et al., 2004). The same calibration strategy has been implemented in published studies (Clark and Donoghue, 2017; Wang and Luo, 2021).

Node_8: Crown Spermatophyta group (Total Angiosperms group)

Minimum age: 0.308 Gya

Maximum age: 0.509 Gya

Justification: Cordaitales is an extinct order of gymnosperms. Morphological fossils of *Cordaixylon iowensis* were found in the middle Pennsylvanian coal-ball locality at What Cheer, Iowa (Trivett, 1992). By following the published study (Su et al., 2021), we constrained the minimum age of the total Angiosperms group at 0.308 Gya based on this fossil record. The maximum age of the total Angiosperms group was set to 0.509 Gya based on the age of the crown Embryophyta group (see Node_7 for details). Note that the "total group of Angiosperms" includes all living members of the crown group of Angiosperms and all their extinct relatives that are more closely related to them than to any other living group (stem group), encompassing the entire evolutionary lineage. Because the fossils used here could be derived from stem group Angiosperms, we can only conservatively place them at the total group Angiosperms, or equivalently, the crown group Spermatophyta, as used in previous studies (Betts et al., 2018; Wang and Luo, 2021).

Node_9: Crown Angiosperm group (Total Eudicots group)

Minimum Age: 0.125 Gya

Maximum Age: 0.25 Gya

Justification: The tricopolate pollen fossil identified from the Cowleaze Chine Member of the Vectis Formation was dated back to 125±1.0 Mya, and was considered as the most ancient evidence of angiosperms (Gradstein et al., 2004). By following the published studies (Clarke et al., 2011; Wang and Luo, 2021), we constrained the minimum age of the crown Angiosperm group at 0.125 Gya.

The maximum age of the crown Angiosperm group was based on the sediments where angiosperm-like pollen has never been found below their first report in the Middle Triassic, corresponding to 247.1±0.2 Mya (Gradstein et al., 2012). Consequently, we constrained the maximum age of the crown Angiosperm group at 0.250 Gya by following the published studies (Clarke et al., 2011; Wang and Luo, 2021) .

Node_10: Total diatom group

Minimum age: 0.14 Gya

Maximum age: 1.891 Gya

Justification: The well-preserved Cretaceous diatom assemblages in Antarctica was dated back to 140 Mya and was considered as the oldest fossil of centric diatom (Harwood et al., 2007). By following the published study (Parfrey et al., 2011; Delaye et al., 2016), we constrained the minimum age of the total diatom group at 0.14 Gya. The maximum age of the total diatom group was set to 1.891 Gya based on the earliest record of simple eukaryote (see Node_5 for details).

Node_11: Crown diatom group (Total pennate diatom group)

Minimum age: 0.08 Gya

Maximum age: 1.891 Gya

Justification: By following the published studies (Parfrey et al., 2011; Delaye et al., 2016), we constrained the minimum age of the crown diatom group (i.e. the total pennate diatom) at 0.08 Gya based on the oldest pennate diatom fossil identified in Carbonaceous cherts of the Tarahumara Formation, Mexico (Chacón-Baca et al., 2002). The maximum age of the crown diatom group was set to 1.891 Gya based on the earliest record of simple eukaryote (see Node_5 for details).

Node_12: Crown Rhodophyta group

Minimum Age: 1.047 Gya, 1.2 Gya, and 1.6 Gya

Maximum Age: 1.891 Gya

Justification: *Bangiomorpha pubescens* was identified in the multicellular fossils in Arctic Canada based on its developmental characters and the distinct shape of cell arrangement. Since these characters are present in other red algae lineages (Yang et al., 2016), we constrained the minimum age of the crown Rhodophyta group by following a recent study (Parfrey et al., 2011) based on the age of the associated Neoproterozoic strata at 1.2 Gya (Butterfield, 2015), or the age of the sedimentary rocks in Baffin Island at 1.047 Gya (Gibson et al. 2018). The fossils of *Rafatazmia chitrakootia* and *Ramathallus lobatus* identified in Chitrakoot Formation were considered to belong to crown-group red algae due to their distinctive morphological features (Bengtson et al., 2017). Although such inference was questioned in later studies (Betts et al. 2018; Gibson et al. 2018; Mills et al. 2022), we constrained the crown Rhodophyta at 1.6 Gya by following the recent study (Strassert et al., 2021b). The maximum age of the crown Rhodophyta group was set to 1.891 Gya based on the earliest record of simple eukaryote (see Node_5 for details).

Node_13: Total Florideophyceae group

Minimum Age: 0.55 Gya

Maximum Age: 1.891 Gya

Justification: The multicellular algae deposited in the phosphatic sediment in Doushantuo Formation of Weng'an, South China, was considered as the lineage that branched near the base of the florideophyte tree (Xiao et al., 2004). Since the age of Doushantuo fossil was estimated to between 0.6 Gya and 0.55 Gya (Knoll, 1999), we constrained the minimum age of the total Florideophyceae group at 0.55 Gya by following the published studies (Parfrey et al., 2011;

Wang and Luo, 2021). The maximum age of the total Florideophyceae group was set to 1.891 Gya based on the earliest record of simple eukaryote (see Node_5 for details).

A5) MCMCTree implementation

Our molecular dating analyses were performed with MCMCTree following the command-line steps detailed in the chapter (Zhang et al., 2021). Since the molecular data are solely composed of protein-coding genes, we used the program CODEML v4.10 to estimate the substitution rates based on the amino acid sequences. The mean substitution rate of the input gene families was then used to inform the Dirichlet-gamma prior (rgene_gamma) to be used in MCMCTree analysis (Reis and Yang, 2011). We then employed a uniform time prior distribution (BDparas=1 1 0) in our molecular dating analysis with MCMCTree due to the little prior knowledge about it, as used in previous studies (dos Reis et al., 2015; dos Reis et al., 2018). To ensure the convergence of MCMC sampling to a posterior distribution, we set the parameter "nsample" and "samplefreq" to "10,000" and "50", respectively, to collect a total of 500,000 samples for each run of the dating analysis. The convergence plot was then made from the repeated runs of each analysis (Fig. S8).

<u>B) Reconstruction of gene gains and losses</u>

The genome content evolution of Cyanobacteria was inferred based on a total of 4,689 orthologous gene families identified in our recent study (Zhang et al., 2021) using gene tree-species tree reconciliation approach. To find out the best-fit reconciliation tools for our dataset, we employed a simulation-based benchmarking analysis by following our recent study [see the section "Inferring gene gain and loss events along species phylogeny" and Fig. S4 in the

Supplementary Information of (Zhang et al., 2023)] and found AnGST and GeneRax as the best-fit MP-based and ML-based reconciliation tools, respectively. We then reconstructed the evolutionary paths of the complete set of orthologous gene family and recovered 37 and 364 gene gains and losses with AnGST and 113 and 342 gains and losses with GeneRax, respectively, at the last common ancestor (LCA) of *Prochlorococcus* without the use of SAGs (Supplementary Data 3).

The functional annotation of the 4,689 gene families was carried out in our recent study by searching against the KEGG database (release 2017) and subsystem database using BLASTP v2.2.6 (Camacho et al., 2009) and the RAST online platform (Aziz et al., 2008), respectively (Supplementary Data 3). Since it is possible that different gene family members were annotated as different functional categories, we employed the majority rule to assign the KEGG and subsystem functional category to each gene family. To validate the genomic changes of important functional genes when *Prochlorococcus* SAGs were included, we performed another round of reconstructions based on the subset of gene family by using the same gene tree-species tree reconciliation approach.

C) Assessing the efficiency of natural selection in deep time

Using the *Synechococcus* clade 5.2 as the reference group, we employed the software RCCalculator (http://www.geneorder.org/RCCalculator/) (Luo et al., 2017) to compare the $d_R/d_C$ ratios (i.e., the relative rate of radical versus conservative nonsynonymous substitutions) of the phylogenetic groups of *Prochlorococcus* (target group) and *Synechococcus* clade 5.1 (control group). A total of 337 single-copy orthologous gene families shared by *Synechococcus* clade 5.1/5.2 and *Prochlorococcus* with SAGs were retrieved from the output of OrthoFinder v2.2.1

(Emms and Kelly, 2015) by allowing the absence of at most five genomes (the same number with SAGs) in a single gene family. Genes were aligned at the amino acid level using MAFFT v7.271 (Katoh and Standley, 2013). DNA sequences were then imposed on these alignments followed by removing the gaps and codons with ambiguous nucleotides. The transition/transversion ratio ($t_S/t_V$) of each gene family was estimated using MEGA-CC v7.0.26 (Kumar et al., 2012) to feed the software RCCalculator. For each gene family, a total of six cases were considered for the calculation of $d_R$ and $d_C$, including two ways of categorizing amino acids (by charge and by volume and polarity) and three approaches of GC-bias correction (uncorrected, on codon frequency correction, and on amino acid composition correction). We then performed the paired *t* test to compare the $d_R/d_C$ ratio between "target" and "control" groups.

**Fig. S1** Diagrams showing the eukaryotic tree topology used in the first-step sequential Bayesian dating analysis, the posterior ages of eukaryotes estimated in the first-step sequential Bayesian dating analysis, and the evolutionary timeline of Cyanobacteria estimated with the focal strategy in the second-step sequential Bayesian dating analysis. (a) the species tree topology used in the first-step dating analysis. Topo-1: the Glaucophyta-basal phylogenomic tree topology in which Prasinodermophyta locates at the base of Viridiplantae. Topo-2: alternative species tree topology in which Rhodophyta group instead of Glaucophyta group positioned at the base of the tree. Topo-3: alternative species tree topology in which Prasinodermophyta locates at the base of Chlorophyta instead of the base of Viridiplantae. Calibrated nodes are marked with solid red circle (see calibration justification in Supplementary Methods). The calibrated nodes “Node_8”, “Node_10” and “Node_11” were not marked due to the absence of the corresponding nodes. These three calibrated nodes were used in the second-step dating analysis. (b) the chronograms of eukaryotic evolution based on Topo-1 and 1-, 3-, 5- , 7-, and 9-partition molecular data. The Akaike Information Criterion (AIC) value for each partition was assessed by IQ-TREE under the auto-determined amino acid substitution model. The posterior ages of the ancestral nodes (start with “Node_e”) were approximated to one of the skew-normal, skew-t and Gamma distributions and used to constrain the time prior of the corresponding nodes in the dating analysis with Cyanobacteria. The vertical dashed line connects the posterior mean age of the crown eukaryotic group. (c) The chronogram of Cyanobacteria evolution estimated with the focal dating strategy. Calibrated nodes are marked with solid circle with specific IDs, which are in consistency with that justified in Supplementary Methods. Note that the nodes start with “Node_e” were constrained with the distributions that were approximated from the eukaryotic dating analysis in panel (b). The vertical bars with green, orange and blue colors represent the time of Tonian, the time of NOE, and the time of Sturtian (left) and Marinoan (right) glaciation, respectively.

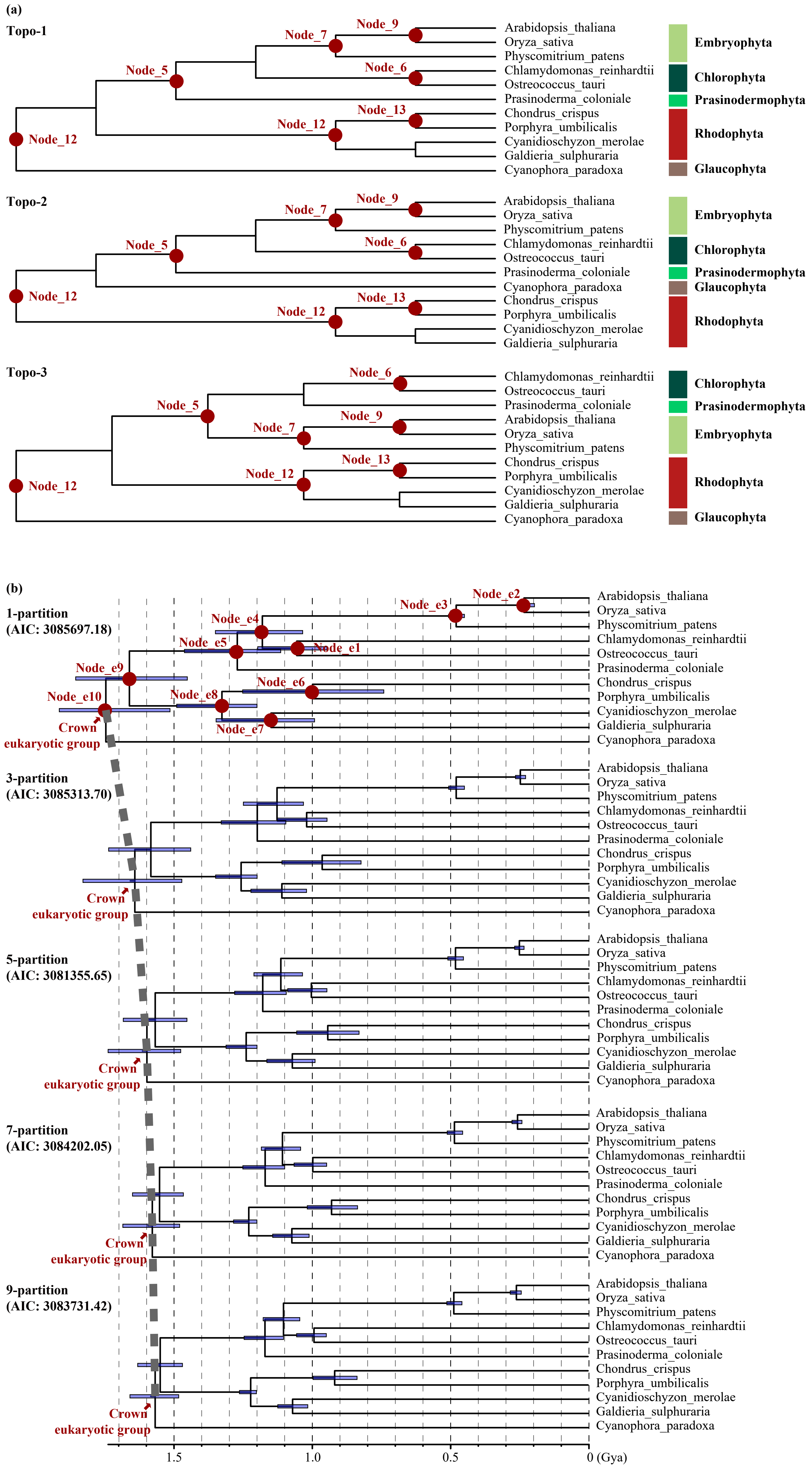

(c)

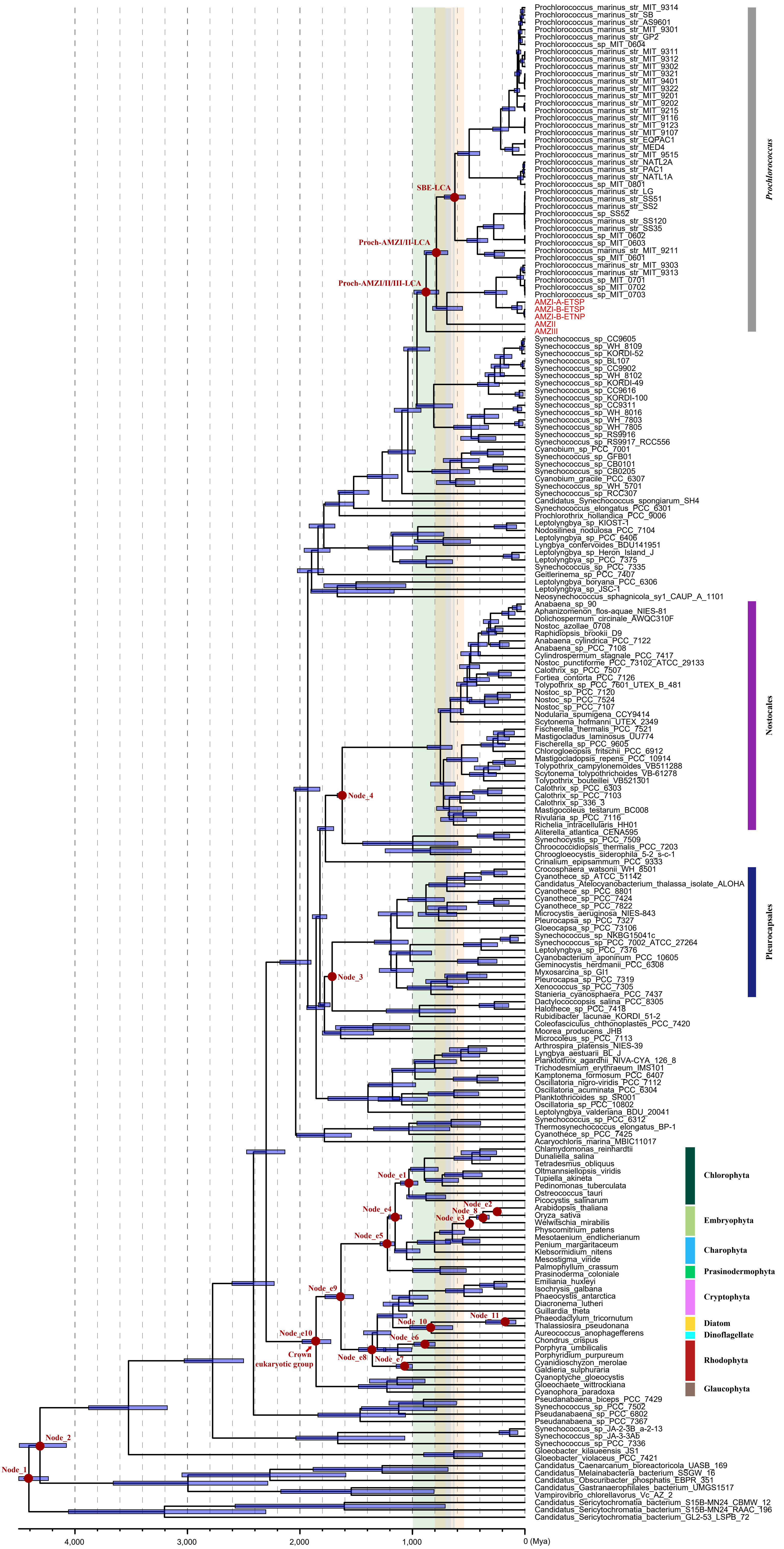

Prochlorococcus_marinus_str_MIT_9314
Prochlorococcus_marinus_str_SB
Prochlorococcus_marinus_str_AS9601
Prochlorococcus_marinus_str_MIT_9301
Prochlorococcus_marinus_str_GP2
Prochlorococcus_sp_MIT_0604
Prochlorococcus_marinus_str_MIT_9311
Prochlorococcus_marinus_str_MIT_9312
Prochlorococcus_marinus_str_MIT_9302
Prochlorococcus_marinus_str_MIT_9321
Prochlorococcus_marinus_str_MIT_9401
Prochlorococcus_marinus_str_MIT_9322
Prochlorococcus_marinus_str_MIT_9201
Prochlorococcus_marinus_str_MIT_9202
Prochlorococcus_marinus_str_MIT_9215
Prochlorococcus_marinus_str_MIT_9116
Prochlorococcus_marinus_str_MIT_9123
Prochlorococcus_marinus_str_MIT_9107
Prochlorococcus_marinus_str_EQPAC1
Prochlorococcus_marinus_str_MED4
Prochlorococcus_marinus_str_MIT_9515
Prochlorococcus_marinus_str_NATL2A
Prochlorococcus_marinus_str_PAC1
Prochlorococcus_marinus_str_NATL1A
Prochlorococcus_sp_MIT_0801
Prochlorococcus_marinus_str_LG
Prochlorococcus_marinus_str_SS51
Prochlorococcus_marinus_str_SS2
Prochlorococcus_sp_SS52
Prochlorococcus_marinus_str_SS120
Prochlorococcus_marinus_str_SS35
Prochlorococcus_sp_MIT_0602
Prochlorococcus_sp_MIT_0603
Prochlorococcus_marinus_str_MIT_9211
Prochlorococcus_sp_MIT_0601
Prochlorococcus_marinus_str_MIT_9303
Prochlorococcus_marinus_str_MIT_9313
Prochlorococcus_sp_MIT_0701
Prochlorococcus_sp_MIT_0702
Prochlorococcus_sp_MIT_0703
AMZI-A-ETSP
AMZI-B-ETSP
AMZI-B-ETNP
AMZII
AMZIII
Synechococcus_sp_CC9605
Synechococcus_sp_WH_8109
Synechococcus_sp_KORDI-52
Synechococcus_sp_BL107
Synechococcus_sp_CC9902
Synechococcus_sp_WH_8102
Synechococcus_sp_KORDI-49
Synechococcus_sp_CC9616
Synechococcus_sp_KORDI-100
Synechococcus_sp_CC9311
Synechococcus_sp_WH_8016
Synechococcus_sp_WH_7803
Synechococcus_sp_WH_7805
Synechococcus_sp_RS9916
Synechococcus_sp_RS9917_RCC556
Cyanobium_sp_PCC_7001
Synechococcus_sp_GFB01
Synechococcus_sp_CB0101
Synechococcus_sp_CB0205
Cyanobium_gracile_PCC_6307
Synechococcus_sp_WH_5701
Synechococcus_sp_RCC307
Candidatus_Synechococcus_spongiarum_SH4
Synechococcus_elongatus_PCC_6301
Prochlorothrix_hollandica_PCC_9006
Leptolyngbya_sp_KIOST-1
Nodosilinea_nodulosa_PCC_7104
Leptolyngbya_sp_PCC_6406
Lyngbya_confervoides_BDU141951
Leptolyngbya_sp_Heron_Island_J
Leptolyngbya_sp_PCC_7375
Synechococcus_sp_PCC_7335
Geitlerinema_sp_PCC_7407
Leptolyngbya_boryana_PCC_6306
Leptolyngbya_sp_JSC-1
Neosynechococcus_sphagnicola_sy1_CAUP_A_1101
Anabaena_sp_90
Aphanizomenon_flos-aquae_NIES-81
Dolichospermum_circinale_AWQC310F
Nostoc_azollae_0708
Raphidiopsis_brookii_D9
Anabaena_cylindrica_PCC_7122
Anabaena_sp_PCC_7108
Cylindrospermum_stagnale_PCC_7417
Nostoc_punctiforme_PCC_73102_ATCC_29133
Calothrix_sp_PCC_7507
Fortiea_contorta_PCC_7126
Tolypothrix_sp_PCC_7601_UTEX_B_481
Nostoc_sp_PCC_7120
Nostoc_sp_PCC_7524
Nostoc_sp_PCC_7107
Nodularia_spumigena_CCY9414
Scytonema_hofmanni_UTEX_2349
Fischerella_thermalis_PCC_7521
Mastigocladus_laminosus_UU774
Fischerella_sp_PCC_9605
Chlorogloeopsis_fritschii_PCC_6912
Mastigocladopsis_repens_PCC_10914
Tolypothrix_campylonemoides_VB511288
Scytonema_tolypothrichoides_VB-61278
Tolypothrix_bouteillei_VB521301
Calothrix_sp_PCC_6303
Calothrix_sp_PCC_7103
Calothrix_sp_336_3
Mastigocoleus_testarum_BC008
Rivularia_sp_PCC_7116
Richelia_intracellularis_HH01
Aliterella_atlantica_CENA595
Synechocystis_sp_PCC_7509
Chroococcidiopsis_thermalis_PCC_7203
Chroogloeocystis_siderophila_5-2_s-c-1
Crinalium_epipsammum_PCC_9333
Crocosphaera_watsonii_WH_8501
Cyanothece_sp_ATCC_51142
Candidatus_Atelocyanobacterium_thalassa_isolate_ALOHA
Cyanothece_sp_PCC_8801
Cyanothece_sp_PCC_7424
Cyanothece_sp_PCC_7822
Microcystis_aeruginosa_NIES-843
Pleurocapsa_sp_PCC_7327
Gloeocapsa_sp_PCC_73106
Synechococcus_sp_NKBG15041c
Synechococcus_sp_PCC_7002_ATCC_27264
Leptolyngbya_sp_PCC_7376
Cyanobacterium_aponinum_PCC_10605
Geminocystis_herdmanii_PCC_6308
Myxosarcina_sp_GI1
Pleurocapsa_sp_PCC_7319
Xenococcus_sp_PCC_7305
Stanieria_cyanosphaera_PCC_7437
Dactylococcopsis_salina_PCC_8305
Halothece_sp_PCC_7418
Rubidibacter_lacunae_KORDI_51-2
Coleofasciculus_chthonoplastes_PCC_7420
Moorea_producens_JHB
Microcoleus_sp_PCC_7113
Arthrospira_platensis_NIES-39
Lyngbya_aestuarii_BL_J
Planktothrix_agardhii_NIVA-CYA_126_8
Trichodesmium_erythraeum_IMS101
Kamptonema_formosum_PCC_6407
Oscillatoria_nigro-viridis_PCC_7112
Oscillatoria_acuminata_PCC_6304
Planktothricoides_sp_SR001
Oscillatoria_sp_PCC_10802
Leptolyngbya_valderiana_BDU_20041
Synechococcus_sp_PCC_6312
Thermosynechococcus_elongatus_BP-1
Cyanothece_sp_PCC_7425
Acaryochloris_marina_MBIC11017
Chlamydomonas_reinhardtii
Dunaliella_salina
Tetradesmus_obliquus
Oltmansiellopsis_viridis
Tupiella_akineta
Pedinomonas_tuberculata
Ostreococcus_tauri
Picocystis_salinarum
Arabidopsis_thaliana
Oryza_sativa
Welwitschia_mirabilis
Physcomitrium_patens
Mesotaenium_endlicherianum
Penium_margaritaceum
Klebsormidium_nitens
Mesostigma_viride
Palmophyllum_crassum
Prasinoderma_coloniale
Emiliania_huxleyi
Isochrysis_galbana
Phaeocystis_antarctica
Diacronema_lutheri
Guillardia_theta
Phaeodactylum_tricornutum
Thalassiosira_pseudonana
Aureococcus_anophagefferens
Chondrus_crispus
Porphyra_umbilicalis
Porphyridium_purpureum
Cyanidioschyzon_merolae
Galdieria_sulphuraria
Cyanoptyche_gloeocystis
Gloeochaete_wittrockiana
Cyanophora_paradoxa
Pseudanabaena_biceps_PCC_7429
Synechococcus_sp_PCC_7502
Pseudanabaena_sp_PCC_6802
Pseudanabaena_sp_PCC_7367
Synechococcus_sp_JA-2-3B_a-2-13
Synechococcus_sp_JA-3-3Ab
Synechococcus_sp_PCC_7336
Gloeobacter_kilaueensis_JS1
Gloeobacter_violaceus_PCC_7421
Candidatus_Caenarcanum_bioreactoricola_UASB_169
Candidatus_Melainabacteria_bacterium_SSGW_16
Candidatus_Obscuribacter_phosphatis_EBPR_351
Candidatus_Gastranaerophilales_bacterium_UMGS1517
Vampirovibrio_chlorellavorus_Vc_AZ_2
Candidatus_Sericytochromatia_bacterium_S15B-MN24_CBMW_12
Candidatus_Sericytochromatia_bacterium_S15B-MN24_RAAC_196
Candidatus_Sericytochromatia_bacterium_GL2-53_LSPB_72
Prochlorococcus
Nostocales
Pleurocapsales
Chlorophyta
Embryophyta
Charophyta
Prasinodermophyta
Cryptophyta
Diatom
Dinoflagellate
Rhodophyta
Glaucophyta
SBE-LCA
Proch-AMZI/II-LCA
Proch-AMZI/II/III-LCA
Node_4
Node_3
Node_e1
Node_e2
Node_8
Node_e3
Node_e4
Node_e5
Node_e9
Node_e10
Crown eukaryotic group
Node_10
Node_11
Node_e6
Node_e8
Node_e7
Node_2
Node_1
4,000
3,000
2,000
1,000
0 (Mya)

**Fig. S2**

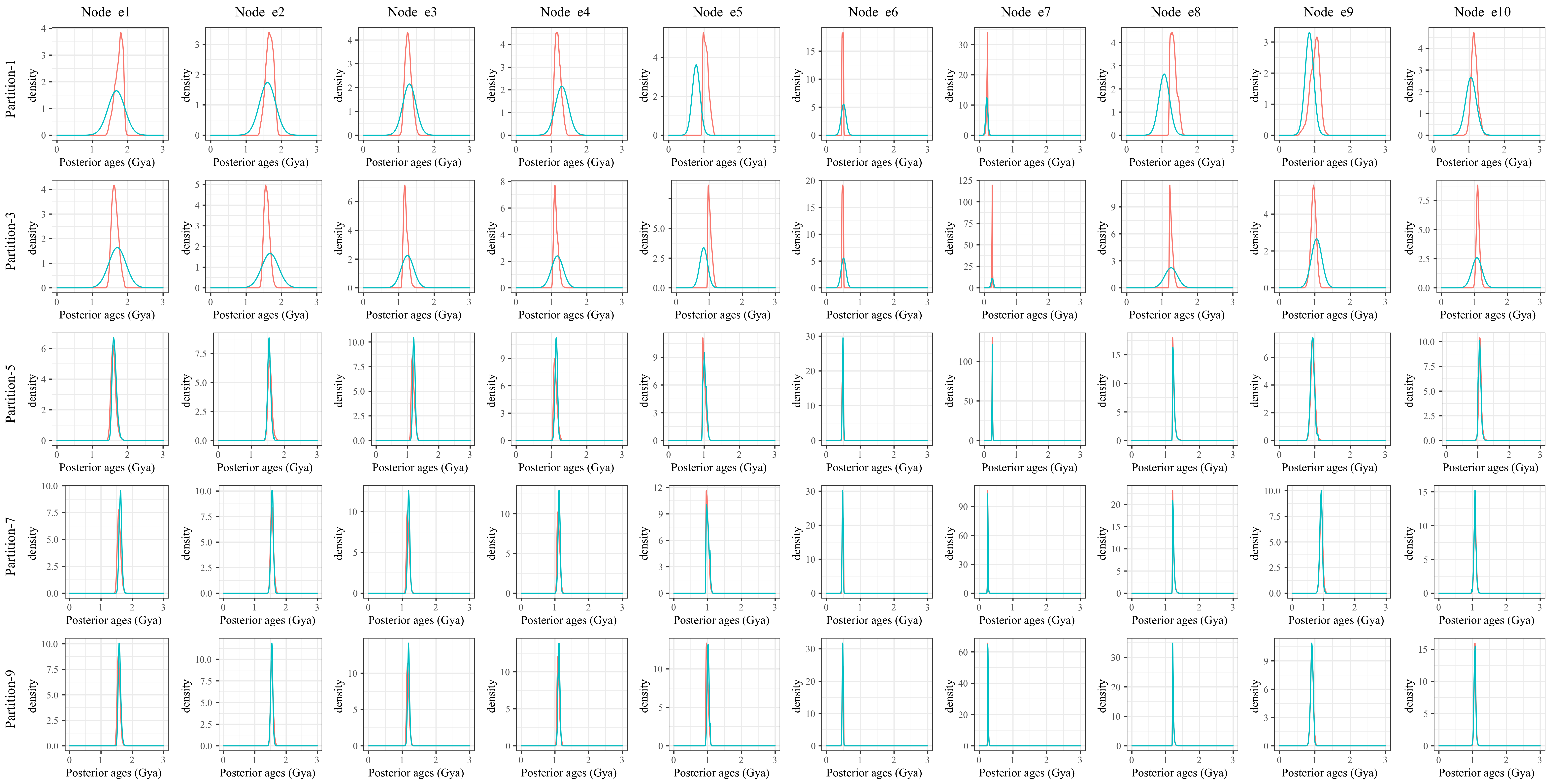


Fig. S2 The plots comparing the posterior ages in the first-step sequential dating analysis (marked with cyan) to the effective time prior in the second-step sequential dating analysis (marked with pink). Large overlaps between them, as in the case of the analyses with 5-, 7- and 9-partition data, suggest that the sequential Bayesian dating approach works well for these analyses. The labels of ancestral nodes (Node_e1 to Node_e10) are consistent with those marked in Fig. S1.

**Fig. S3**

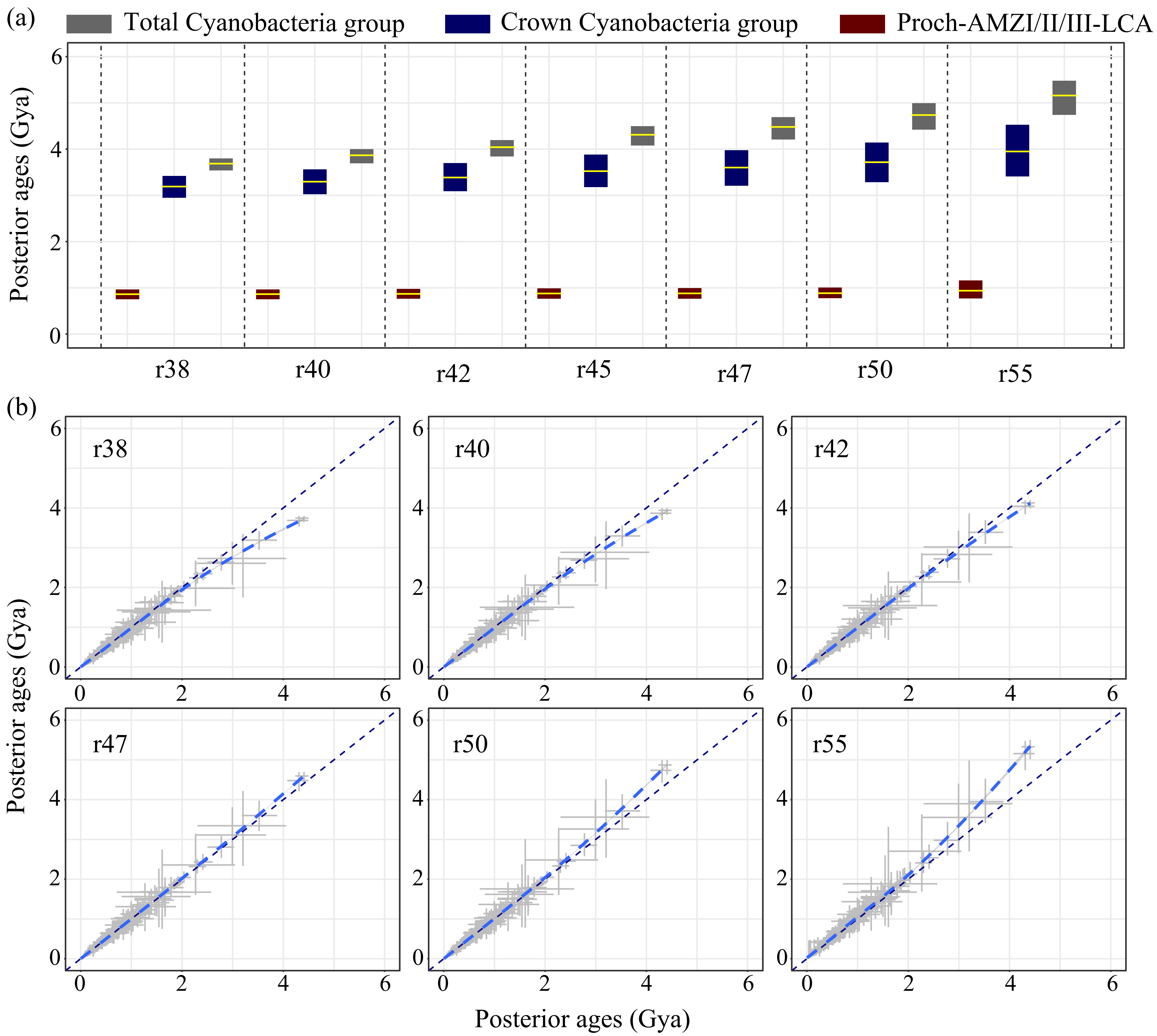


Fig. S3 The posterior ages that estimated with different root maximum ages. (a) The posterior ages of Cyanobacteria and *Prochlorococcus*. The horizontal lines and the vertical bars represent the mean and the 95% highest probability density (HPD) intervals of posterior age estimates. (b) The posterior ages estimated with the root maximum age at 4.5 Gya (x-axis) and that estimated with alternative root maximum ages (y-axis). The blue curves are trend lines estimated using the “loess” method in the R package “ggplot2”. In both panels, the root maximum ages at 3.8 Gya, 4.0 Gya, 4.2 Gya, 4.5 Gya, 4.7 Gya, 5.0 Gya and 5.5 Gya were marked with “r38”, “r40”, “r42”, “r45”, “r47”, “r50” and “r55”.

**Fig. S4**

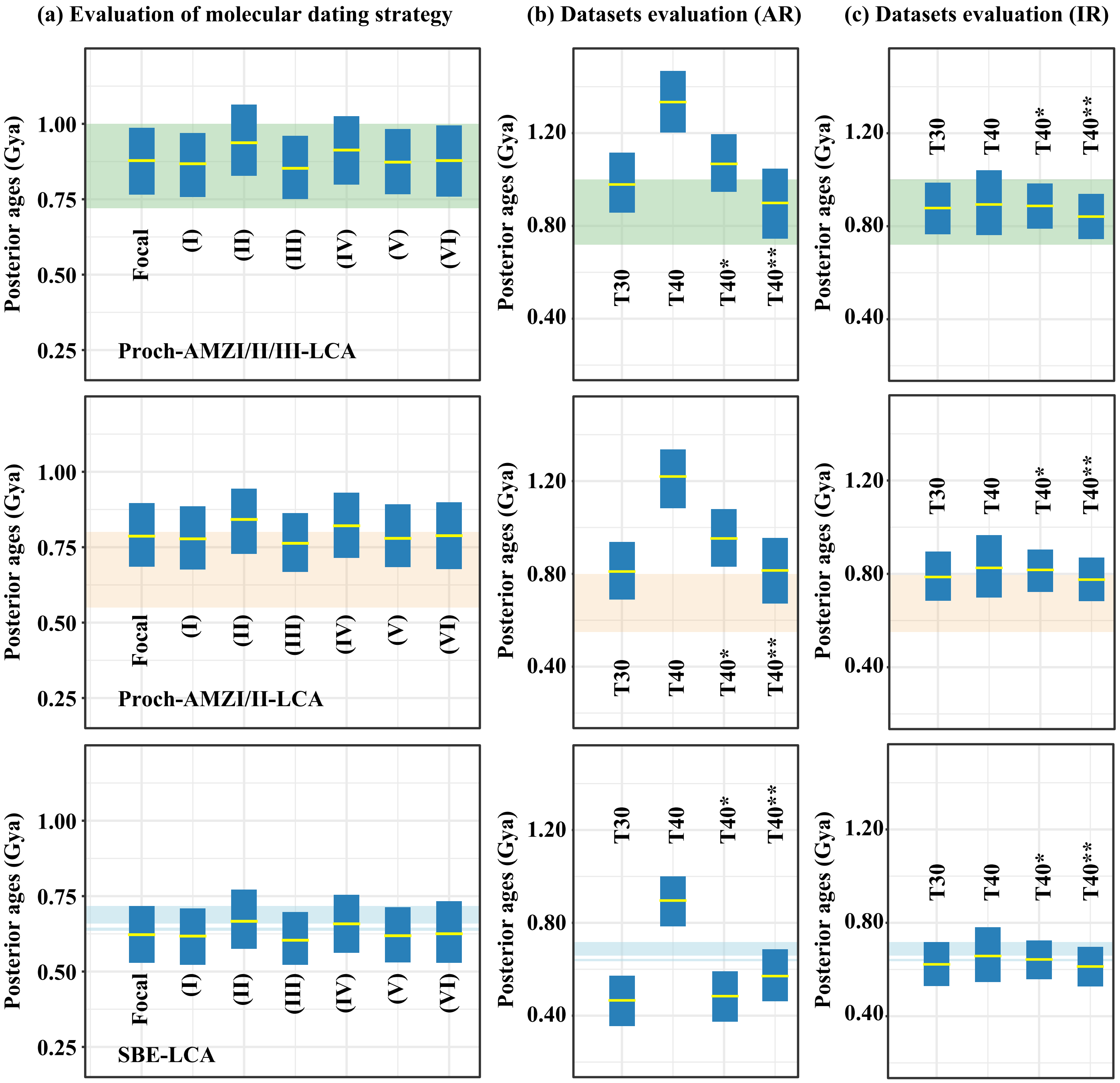


Fig. S4 Posterior ages of the three ancestral nodes Proch-AMZI/II/III-LCA, Proch-AMZI/II-LCA and SBE-LCA estimated with different dating strategies. (a) Dating analysis with the dataset "T30" and the optimal clock model (IR model) under focal and alternative strategies. Focal strategy: the Glaucophyta-basal phylogenomic tree was used, in which Prasinodermophyta locates at the base of Viridiplantae. The minimum age constraints for the total Nostocales group and the crown Rhodophyta group are 1.6 Gya and 1.2 Gya, respectively. I and II, alternative minimum age constraint of the total Nostocales group at 1.2 Gya and 2.0 Gya, respectively; III and IV, alternative minimum age constraint of the crown Rhodophyta group at 1.047 Gya and 1.6 Gya, respectively; V: alternative species tree topology in which Rhodophyta instead of Glaucophyta represent the earliest-split branch of Eukaryotes; VI: alternative species tree topology in which Prasinodermophyta locates at the base of Chlorophyta instead of the base of Viridiplantae. (b) and (c) The posterior ages of the Proch-AMZI/II/III-LCA, Proch-AMZI/II-LCA and SBE-LCA estimated with different molecular datasets under auto-correlated rate (AR) model and independent rate (IR) model, respectively (see details in Supplementary Methods). The yellow lines and the blue vertical bars represent the mean and the 95% highest probability density (HPD) intervals of posterior age estimates. The green, orange, and blue horizontal bars in panels with "Proch-AMZI/II/III-LCA", "Proch-AMZI/II-LCA" and "SBE-LCA" represent the time of Tonian, NOE, and Sturtian (upper) and Marinoan (lower) glaciation, respectively. Note that these glaciation events are expected to have occurred on the branch leading to the ancestral node SBE-LCA, but the panel "SBE-LCA" shows the estimated age of the node SBE-LCA.

Fig. S5

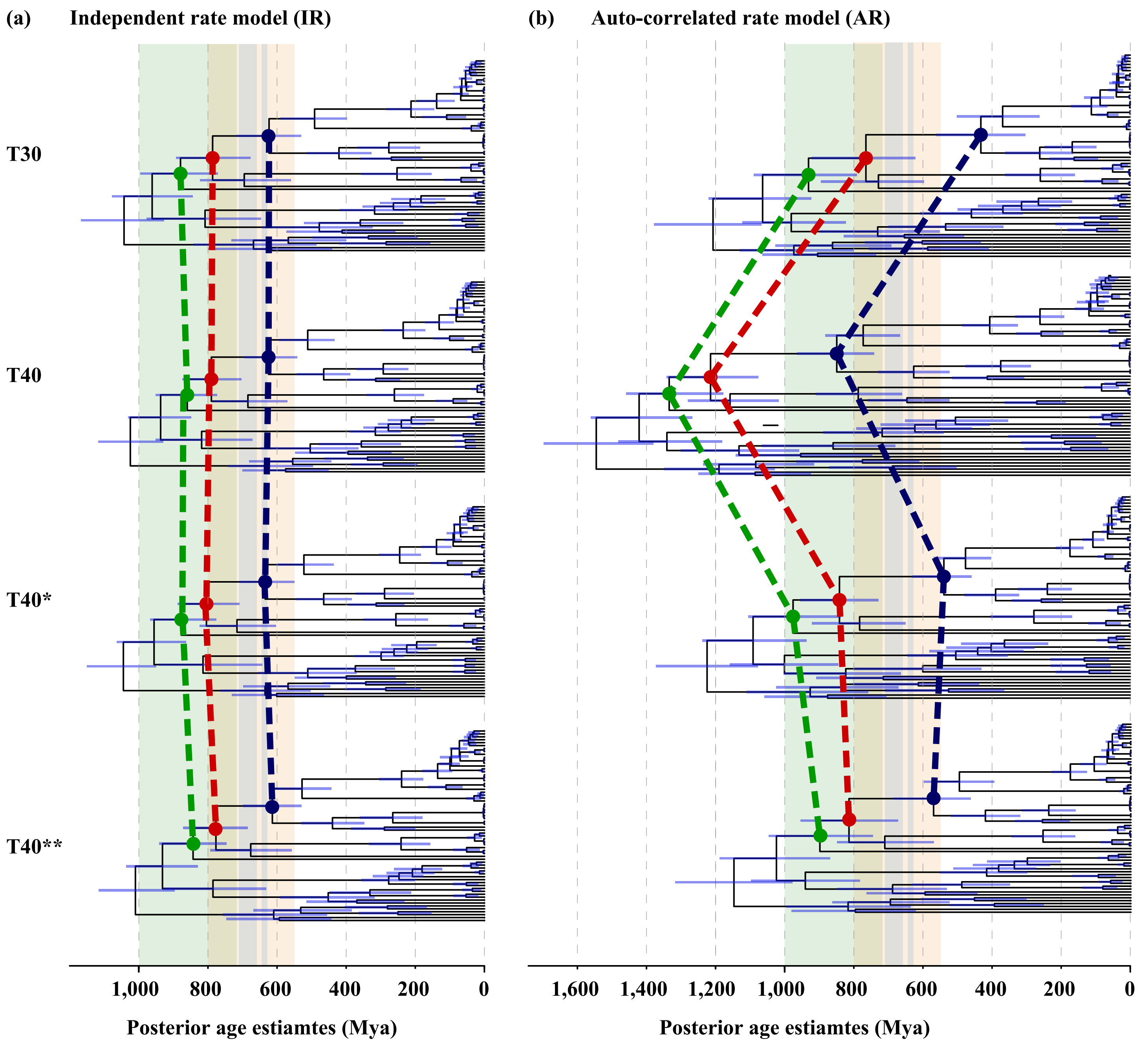


Fig. S5 The chronogram showing the variations of posterior age estimates by using different datasets under independent rate (IR) model or auto-correlated (AR) model. The green, red and blue dashed lines connect to Proch-AMZI/II/III-LCA, Proch-AMZI/II-LCA and SBE-LCA, respectively. The green, orange and blue vertical bars in both panels represent the time of Tonian, the time of NOE and the time of Sturtian (left) and Marinoan (right) glaciation, respectively. The horizontal blue bars on ancestral nodes represent the 95% highest probability density (HPD) intervals of posterior age estimates. For all the analyses except those with the dataset “T40” under the AR model, we found that the branch leading to the SBE-LCA encompasses the Snowball Earth glaciations.

Fig. S6

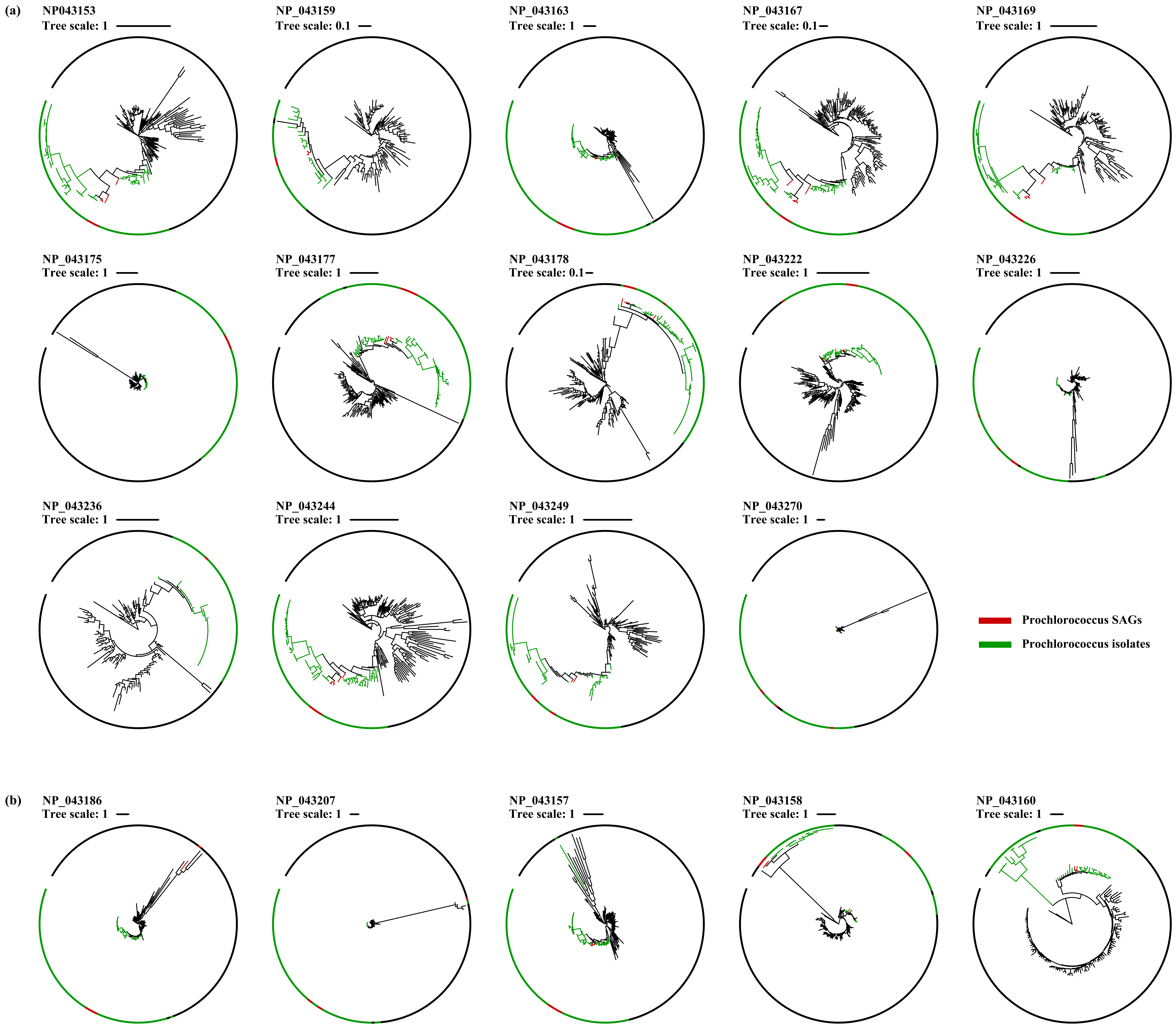


Fig. S6 The maximum likelihood phylogeny of the gene families involved in the dataset "T40". (a) The phylogenetic trees of the 14 gene families in which *Prochlorococcus* forms a monophyletic group. (b) The phylogenetic trees of five gene families in which *Prochlorococcus* forms paraphyletic or polyphyletic groups. *Prochlorococcus* isolates and SAGs are marked with green and red, respectively.

**Fig. S7**

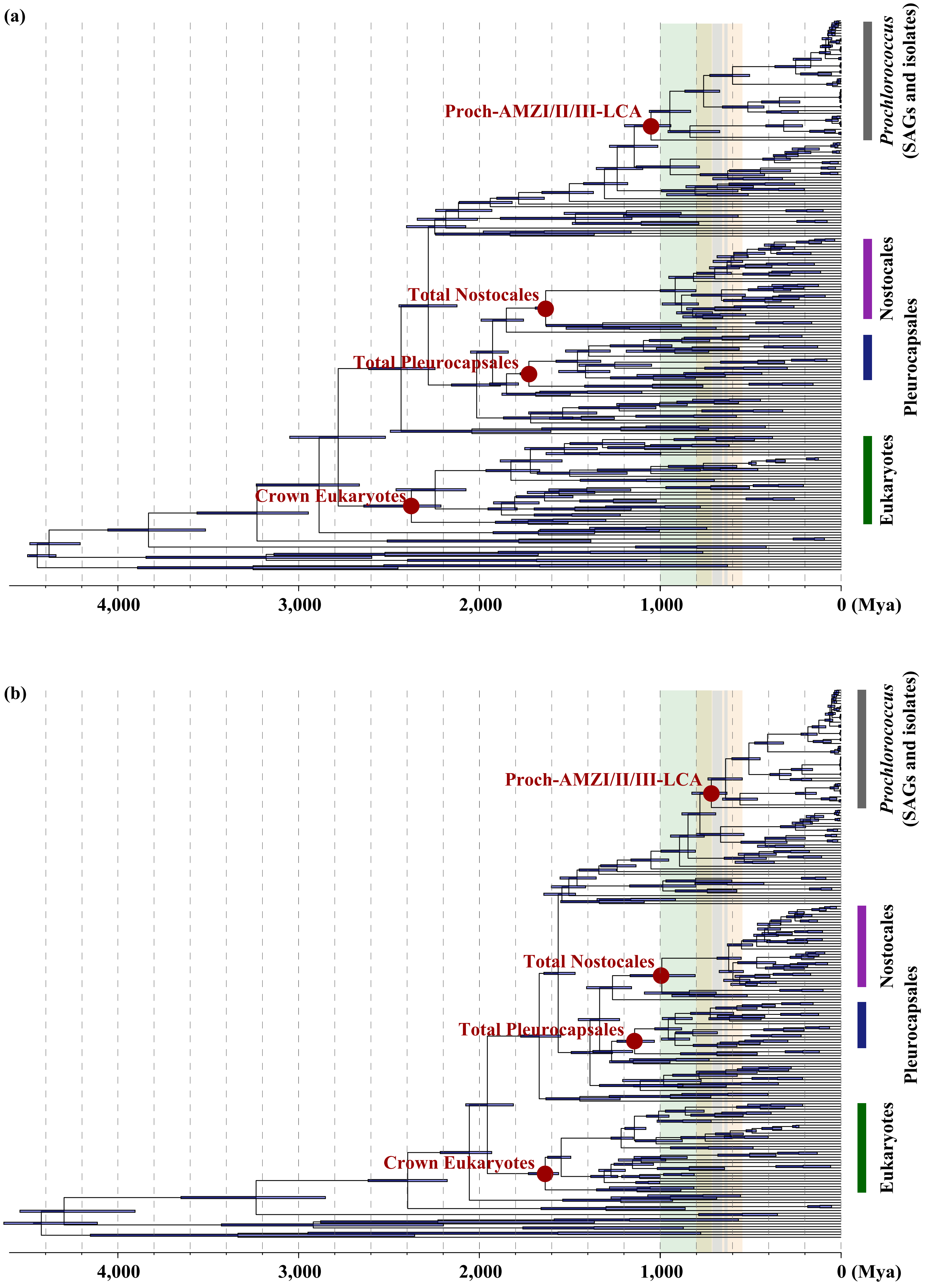


Fig. S7 The chronogram of *Prochlorococcus* estimated with different dating strategies. (a) The chronogram of Cyanobacteria evolution estimated without using the sequential Bayesian dating method. (b) The chronogram of Cyanobacteria evolution estimated with sequential Bayesian dating method but based on soft bound calibration. In both panels, the green, orange and blue vertical bars represent the time of Tonian, the time of NOE and the time of Sturtian (left) and Marinoan (right) glaciation, respectively.

**Fig. S8**

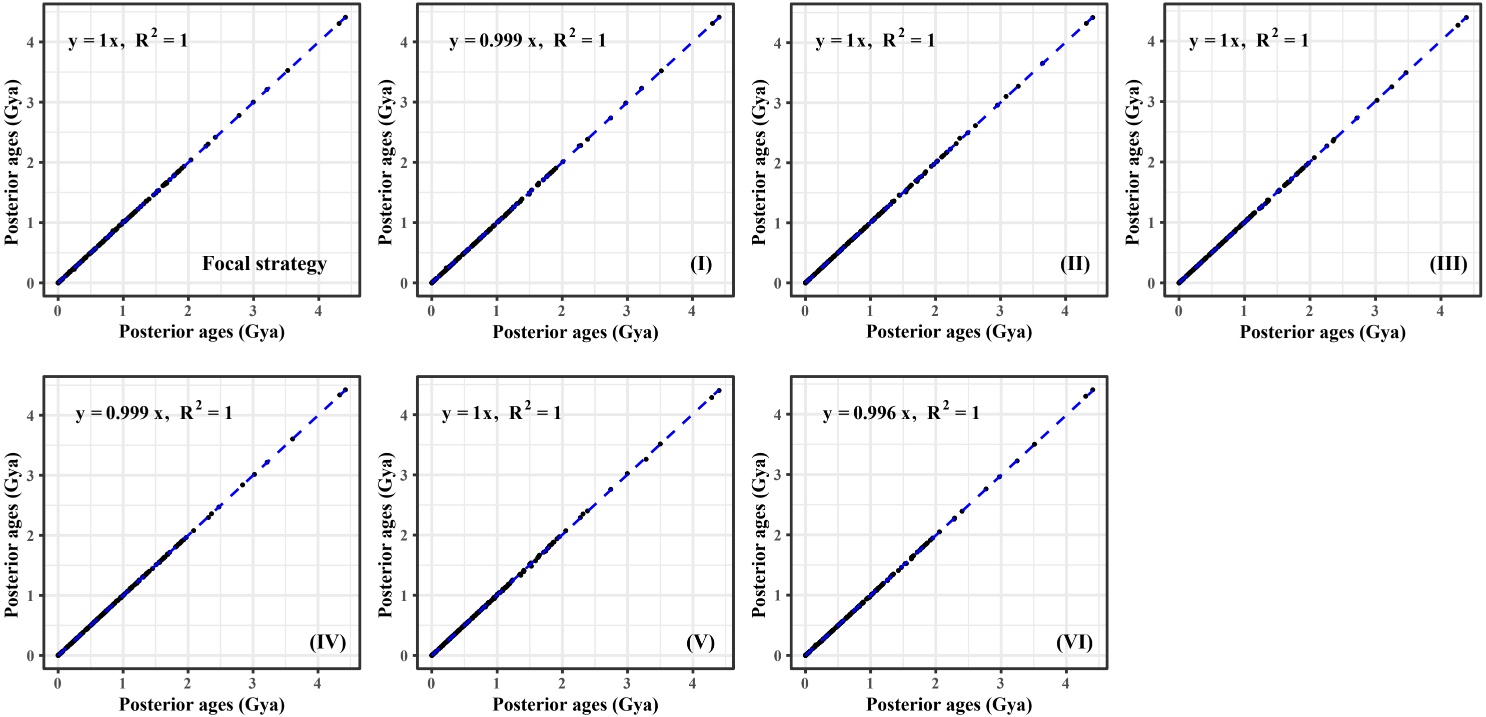


Fig. S8 Correlations of the posterior mean ages between replicated MCMC runs in dating analyses with different strategies. Focal strategy: Glaucophyta-basal phylogenomic tree was used, in which Prasinodermophyta locates at the base of Viridiplantae. The minimum age constraints for the total Nostocales group and the crown Rhodophyta group are 1.6 Gya and 1.2 Gya, respectively. I and II, alternative minimum age constraint of the total Nostocales group at 1.2 Gya and 2.0 Gya, respectively; III and IV, alternative minimum age constraint of the crown Rhodophyta group at 1.047 Gya and 1.6 Gya, respectively; V: alternative species tree topology in which Rhodophyta instead of Glaucophyta represent the earliest-split branch of the Eukaryote; VI: alternative species tree topology in which Prasinodermophyta locates at the base of Chlorophyta instead of the base of Viridiplantae. Partitioned data with different number of categories were used for dating analysis. Convergence of the independent run is achieved if points fall almost perfectly on the y=x line.

**Table S1** The minimum (Min.) and maximum (Max.) age constraints applied in the present study. The "Node_ID" labels are in consistency with that marked in Fig. S1. See calibration justifications in Supplementary Methods.

| Node_ID | Node_label | Min. (Gya) | Max. (Gya) |
|---|---|---|---|
| Node_1 | Root | 2.32 | 4.5 |
| Node_2 | Total oxygenic Cyanobacteria | 2.32 | 4.5 |
| Node_3 | Total group of Pleurocapsales | 1.7 | 4.5 |
| Node_4 | Total Nostocales group | 1.2/1.6/2.0 | 4.5 |
| Node_5 | Crown Viridiplantae group | 0.947 | 1.891 |
| Node_6 | Crown Chlorophyta group | 0.947 | 1.891 |
| Node_7 | Crown Embryophyta group | 0.45 | 0.509 |
| Node_8 | Crown Spermatophyta group (Total Angiosperms group) | 0.308 | 0.509 |
| Node_9 | Crown Angiosperm group (Total Eudicots group) | 0.125 | 0.25 |
| Node_10 | Total diatom group | 0.14 | 1.891 |
| Node_11 | Crown diatom group (Total pennate diatom group) | 0.08 | 1.891 |
| Node_12 | Crown Rhodophyta group | 1.047/1.2/1.6 | 1.891 |
| Node_13 | Total Florideophyceae group | 0.55 | 1.891 |

Table S2 The approximated distributions of posterior ages (Gya) estimated from the Eukaryotic dating analysis based on alternative species tree topologies. ST: skew-t distribution; SN: skew-normal distribution; G: Gamma distribution. Partitioned datasets (P1, P3, P5, P7 and P9) were obtained by using Gaussian Mixture Model (GMM) clustering method. The labels of node ID and topology are in consistency with that marked in Fig. S1.

| Partition | Node_ID | Topo-1 | Topo-2 | Topo-3 |
|---|---|---|---|---|
| P1 | Node_e1 | ST(0.947009028237519,0.132083401278449,188831.374050154,22521.7393320934) | SN(1.03992382158513,0.0701604525492831,0.995271699991409) | ST(0.947004994849179,0.14784423711763,1257687.30099645,754303.420887133) |
| | Node_e2 | ST(0.253928976194357,0.0211295016879279,-5.30705063024386,5.18700815515858) | ST(0.25411535401928,0.0211923280638896,-5.37332378145166,4.5346734528158) | SN(1.15726483538633,0.0959032444211549,0.878186805914551) |
| | Node_e3 | G(760.433540510967,1584.2483470738) | G(751.251689051131,1564.89663714011) | ST(0.253843349599167,0.0238296037789911,-6.21989757626896,4.9322776598979) |
| | Node_e4 | SN(1.17983867004749,0.084760323516396,0.612911313715537) | ST(1.06295607968192,0.122914678154335,3.63209146057269,43.0183717794835) | SN(0.48058054862803,0.0173979755544496,6.72920565240889e-06) |
| | Node_e5 | SN(1.27074998812116,0.0921979572652836,0.51219538245251 8) | ST(1.14352543124248,0.129699829738652,2.84815623871883,77.4200399791712) | SN(1.27206409547933,0.104131883449184,0.709800337948637) |
| | Node_e6 | SN(0.999650045665928,0.132304141486051,-0.506286621685569) | SN(1.52652003193354,0.117263710990939,0.38521844565696) | ST(1.08183775836911,0.144897928666716,-0.663224966896689,20.171414055666) |
| | Node_e7 | SN(1.14922842358081,0.0892528542202028,0.346724181797548) | ST(1.21192149810159,0.210941953294621,-1.75699371244096,48.9315646667928) | ST(1.05934675697091,0.14349817575613,2.13645936584218,46.6890437309504) |
| | Node_e8 | ST(1.20002867408094,0.153559787596852,256001.983202595,93294.1665628673) | SN(1.22301926663068,0.126947169269178,0.62078201688409) | ST(1.2000372195224,0.170449141037837,3091282.82445646,82892.830178321) |
| | Node_e9 | SN(1.66165962032491,0.106691593386558,-0.329045440462549) | SN(1.4062908576564,0.135988773635197,0.926491820130163) | SN(1.65376413413884,0.111658486886544,-0.0557009674653375) |
| | Node_e10 | SN(1.7484265472834,0.113162802143504,-0.899283855050421) | SN(1.69373689236064,0.120530438349803,-0.403207001446451) | SN(1.73751955594834,0.119057102679252,-0.886788886214801) |
| P3 | Node_e1 | ST(0.947008716944399,0.0925672818638039,395053.251854879,5534.54257810165) | ST(0.946998012363959,0.0411252289461505,15691.9250933088,3.45101282854242) | SN(1.01776525248907,0.0534604872388145,0.995271699978451) |
| | Node_e2 | ST(0.251723582525024,0.00664229612320892,-0.698672367120501,3.32987805111226) | ST(0.25184836957222,0.00678659172000364,-0.672431850800888,4.1703273925104) | ST(1.02031650935779,0.0888753722414672,7.21449309996149,16.4726200516829) |
| | Node_e3 | G(804.767007390193,1677.42605556989) | ST(0.45047645334574,0.0335694280279941,54.3458323666912,50.0140444717586) | ST(0.25246078307002,0.00782194429764302,-1.08555486406752,3.81381999384993) |
| | Node_e4 | SN(1.12801130825149,0.0603974279406597,0.872561547837008) | ST(1.03960588909721,0.0617534654237928,4.43990455507023,7.56230741839981) | G(797.407180003467,1661.79321424019) |
| | Node_e5 | SN(1.1995040377637,0.063628471387562,0.833567298926261) | ST(1.10350940586686,0.0680635561620616,3.78318008886796,8.67353350136696) | ST(1.11854172779681,0.0988095374467337,4.85818445389617,23.3208293624188) |
| | Node_e6 | SN(0.964465950013851,0.0723032277032118,-0.124220454416856) | SN(1.46054451301772,0.0804626120409564,0.551833437093586) | ST(1.01375178003583,0.0840658699259954,-0.709565191426236,17.5988155589876) |
| | Node_e7 | ST(1.06318471785065,0.0654388480049158,1.73799537420501,18.4735412427754) | ST(0.943623056598497,0.123810841236681,1.60736048316811,44.5973585328924) | ST(1.05965455534799,0.0760549888266332,2.11362752384224,11.7645750094172) |
| | Node_e8 | ST(1.20004632126876,0.073605063385967,1109.27328598865,107.983487989289) | ST(1.06736341272873,0.135021464775182,4.52092068155965,18.2686564113205) | SN(1.26946596214367,0.0524854352056698,0.995271699997076) |
| | Node_e9 | SN(1.58403764376203,0.0786855590608889,0.511952534189067) | SN(1.33050599245495,0.0973937610222073,0.99481086508932) | SN(1.59856584343085,0.0874266350667241,0.643595183907186) |
| | Node_e10 | SN(1.64188571664032,0.0945532059065799,0.550539633487648) | SN(1.60187388400139,0.0883540487219101,0.734035816539049) | SN(1.65704315823693,0.100426040389428,0.587784489611719) |
| P5 | Node_e1 | ST(0.947001643099423,0.0710541431727492,788381.189942415,212225.442734886) | ST(0.946992125897533,0.0218520618282554,1697.91231866155,3.11258169617859) | ST(0.947007826832104,0.0649466236421879,1003639.94781409,1318257.94811536) |
| | Node_e2 | ST(0.250127148324976,0.00598303580602628,0.0684349617765512,4.04951751856738) | ST(0.248829890790786,0.00630380745836246,0.501659072396194,4.02116230269592) | ST(1.01930393945217,0.0681455779881077,6.74074235156932,22.9437805377277) |
| | Node_e3 | SN(0.481668598720216,0.0168197250190611,-0.0160282515363059) | SN(0.481330316751239,0.0166766468955682,-0.0158436552978234) | ST(0.250198991902348,0.00591515099427599,-0.0353521765714038,3.94737820954197) |
| | Node_e4 | SN(1.1134652990811,0.0478441042749751,0.847571337254799) | ST(1.04117097209899,0.0372844127591281,3.03962670103255,7.43148958373722) | SN(0.484180219097593,0.018566504161019,-0.952617087937479) |
| | Node_e5 | SN(1.17901487233515,0.0502730091846881,0.781683445755314) | ST(1.09887910902477,0.0434040399099312,2.82798728996366,9.40992757141088) | ST(1.11843654596682,0.0758668780067156,4.37962260852658,40.0709044215021) |
| | Node_e6 | ST(0.906097113892492,0.0663860287983683,0.960654209006884,24.8341525147002) | ST(1.35662571880573,0.0883543232746611,2.31933807917909,32.2221213538911) | ST(0.916845095370732,0.0658052627442176,0.630140655315418,76.2974160263401) |
| | Node_e7 | ST(1.03836891307365,0.0483543666758983,1.31434737764259,8.37069169737928) | ST(0.947575694256977,0.0855578058746048,1.12291351095913,16.9794526297728) | ST(1.04345942658838,0.049279102919206,1.26986957974688,8.61258388298607) |
| | Node_e8 | ST(1.20004355618593,0.0385939971265623,163.893406778112,3.8286212521273) | ST(1.05181571491354,0.0983512992542517,3.32115677229495,9.5524720216696) | ST(1.19999957025594,0.0479085170256309,44131.2278165209,5.64009186243413) |
| | Node_e9 | ST(1.51116459557198,0.0787329384305305,1.81685467515153,20.6969761889401) | ST(1.20002201408201,0.130467975076728,49627.0900068496,198281.632562559) | SN(1.58053701235691,0.0635918271331723,0.535647012413674) |
| | Node_e10 | ST(1.52976987216774,0.0931468058152007,2.067533795369,30.7423093644254) | SN(1.57497075613027,0.068892710459673,0.582113180984603) | SN(1.61226512209239,0.0702600907111078,0.549462190156318) |
| P7 | Node_e1 | ST(0.947014131461715,0.0620222217397256,2679913.2033662,1080392.31707031) | ST(0.946998569872503,0.0174732680398857,18045.7082856377,4.11804096603651) | ST(0.947002696769579,0.0406694843412252,5215.84999113271,9.26941638549794) |
| | Node_e2 | ST(0.247104348947452,0.00925492576513621,1.74794876126143,4.76699760096625) | ST(0.247516217462948,0.0104690981200336,2.1498448458078,5.13430539086492) | ST(1.02084357206764,0.0483998492583989,4.86794068046945,21.7401060033167) |
| | Node_e3 | SN(0.484607005849542,0.0177897135822464,-0.935055282405288) | SN(0.484282219571827,0.017333137166264,-0.882175899150407) | ST(0.247607934543968,0.00836822404912364,1.33956444098875,4.71219683050041) |
| | Node_e4 | SN(1.10689403625262,0.0393648231562773,0.767562136421996) | ST(1.04271067177859,0.0294559732018967,2.45383782608548,18.0700002154352) | SN(0.485668221610284,0.0176180988010783,-0.952605048313685) |
| | Node_e5 | SN(1.16977497132218,0.0409439060442821,0.690462405326911) | ST(1.10016939418064,0.0328727816728911,2.10255656440292,23.8414126458638) | ST(1.12033110188286,0.053622431966359,3.23163952853346,28.9445127095576) |
| | Node_e6 | ST(0.912146296006737,0.0462400804452374,0.495591133968037,16.9357172486013) | SN(1.41040712279752,0.0480285122962032,0.338648534607267) | SN(0.932081542540719,0.0486896010342068,0.135447680524629) |
| | Node_e7 | ST(1.04837959059317,0.0382792549961809,1.18800751236539,13.7941479438057) | G(265.279916425779,271.522504988838) | ST(1.05190707640601,0.0390212267930665,1.09645252282407,8.97719828340625) |
| | Node_e8 | ST(1.19999111033445,0.031206063072994,441.506155162772,4.89007189211235) | ST(1.05982803146673,0.0734000733395677,2.7481615161138,12.4059409881786) | ST(1.20000865476948,0.0439902701139929,288.1263295098,232.379912266433) |
| | Node_e9 | SN(1.55294686792331,0.0477670251853834,0.464255971348281) | SN(1.27765996149274,0.0584258504133728,0.995226804246492) | ST(1.50504426563871,0.071669521609028,2.01904350526644,35.5830419920521) |
| | Node_e10 | SN(1.57911577811616,0.0539341368438662,0.51350878448769) | SN(1.54589877428123,0.0514477059513037,0.430983451130804) | ST(1.52354241828757,0.0819231626628049,2.14272212401583,30.0967904045398) |
| P9 | Node_e1 | SN(0.991111894751987,0.03332445635735,0.995271699979887) | ST(0.946999236726186,0.0143290073200169,12413.8731683498,3.1913001457748) | ST(0.947005932334076,0.0458847356758509,382810.216192232,494506.398791883) |
| | Node_e2 | ST(0.247240252744649,0.0145578147255953,3.39427099661804,11.4072786651291) | ST(0.247929632067641,0.0183099105537815,4.19074600564716,39.0016561417067) | ST(1.02286161854232,0.0480862710291324,5.55572189902797,33.7931299327908) |
| | Node_e3 | SN(0.486522845732729,0.0171405972180714,-0.954067161726615) | SN(0.486048788189541,0.0168632876373028,-0.935480432676684) | ST(0.247073592599232,0.0127710657472041,2.98498604636564,7.54796667103066) |
| | Node_e4 | SN(1.10180937577604,0.035154323391378,0.779413872713466) | ST(1.04210416100848,0.026868299682466,2.71345441949559,11.8350421342536) | SN(0.487214690074658,0.0168719645334604,-0.966118370900897) |
| | Node_e5 | SN(1.16899086177198,0.0369679798139521,0.69581515411529) | ST(1.10264983583735,0.0312619608878134,2.31048682820804,18.4617953960039) | ST(1.12117927782902,0.0513307051087936,3.40874994170281,29.7588664673281) |
| | Node_e6 | ST(0.915663606820819,0.0387422758341086,0.0317882114586939,33.9360529019422) | SN(1.40531803210462,0.0438866142590941,0.418749555658464) | ST(0.90158826366547,0.0446191504300749,0.609109776628593,27.8582466250236) |
| | Node_e7 | ST(1.05250834714857,0.0283164613133837,0.912087607063987,9.52169127146137) | SN(0.969847316159917,0.0600260867575324,0.408931911390068) | ST(1.05272698840436,0.0319688429786453,1.10386633201021,11.5941377100326) |
| | Node_e8 | ST(1.20000436339395,0.0203338955685574,4753.67534988378,3.57348940294987) | ST(1.05632538938211,0.0855168788686523,4.18192362202497,20.4986375369667) | ST(1.20000059251832,0.0240056055572981,7835.17061581574,3.77963183776864) |
| | Node_e9 | SN(1.54811242575812,0.0423841401518277,0.478730122191356) | ST(1.20005591998935,0.0992704556565329,1678915.58995703,191547.83296711) | SN(1.55994004167898,0.0454927198351836,0.44413552818217) |
| | Node_e10 | ST(1.5183859866529,0.0657779020382148,2.10179285974435,68.6576553923665) | SN(1.55300855296583,0.0512422659197121,0.558825446227796) | ST(1.53015927326821,0.0685827836788622,1.88492231487419,76.1486868513453) |
